# Histone variants in archaea and the evolution of combinatorial chromatin complexity

**DOI:** 10.1101/2020.04.13.037952

**Authors:** Kathryn M Stevens, Jacob B Swadling, Antoine Hocher, Corinna Bang, Simonetta Gribaldo, Ruth A Schmitz, Tobias Warnecke

## Abstract

Nucleosomes in eukaryotes act as platforms for the dynamic integration of epigenetic information. Post-translational modifications are reversibly added or removed and core histones exchanged for paralogous variants, in concert with changing demands on transcription and genome accessibility. Histones are also common in archaea. Their role in genome regulation, however, and the capacity of individual paralogs to assemble into histone-DNA complexes with distinct properties remain poorly understood. Here, we combine structural modelling with phylogenetic analysis to shed light on archaeal histone paralogs, their evolutionary history and capacity to generate complex combinatorial chromatin states through hetero-oligomeric assembly. Focusing on the human commensal *Methanosphaera stadtmanae* as a model archaeal system, we show that the heteromeric complexes that can be assembled from its seven histone paralogs vary substantially in DNA binding affinity and tetramer stability, occupying a large but densely populated chromatin state space. Using molecular dynamics simulations, we go on to identify unique paralogs in *M. stadtmanae* and *Methanobrevibacter smithii* that are characterized by unstable dimer:dimer interfaces. We propose that these paralogs act as *capstones* that prevent stable tetramer formation and extension into longer oligomers characteristic of model archaeal histones. Importantly, we provide evidence from phylogeny and genome architecture that these capstones, as well as other paralogs in the Methanobacteriales, have been maintained for hundreds of millions of years following ancient duplication events. Taken together, our findings indicate that at least some archaeal histone paralogs have evolved to play distinct and conserved functional roles, reminiscent of eukaryotic histone variants. We conclude that combinatorially complex histone-based chromatin is not restricted to eukaryotes and likely predates their emergence.

## INTRODUCTION

Cells dynamically regulate access to genomic information in response to upstream signals. This may involve wholesale remodelling of chromatin, for example during spermatogenesis where histones are largely replaced by protamines. Other changes in chromatin state are less radical, involving tweaks to pre-existing chromatin architecture. In eukaryotes, the nucleosome provides the principal platform for such tweaks, prominently via post-translational modifications (PTMs) but also through the exchange of core histones for paralogous variants (Henikoff and Smith 2015). Like PTMs, histone variants can alter nucleosome dynamics or affect the recruitment of *trans* factors to reinforce existing chromatin states, establish new ones, or poise chromatin for future change. In many cases, such paralog exchange is regulated and adaptive. For example, in humans, *de novo* deposition of one histone variant (H2A.X) and eviction of another (H2A.Z) facilitate repair of UV-induced double strand breaks (Piquet *et al.* 2018).

Histones are not restricted to eukaryotes but also common in archaea, where they assemble into tetramers that are structurally very similar to the (H3-H4)_2_ tetramers at the core of eukaryotic nucleosomes (Figure 1A) (Decanniere *et al.* 2000; Malik and Henikoff 2003; Mattiroli *et al.* 2017). In some archaea, including the model species *Methanothermus fervidus* and *Thermococcus kodakarensis*, additional histone dimers can be tagged onto this tetramer to yield oligomers of increasing length that wrap consecutively more DNA (Maruyama *et al.* 2013; Nalabothula *et al.* 2013; Mattiroli *et al.* 2017; Henneman *et al.* 2018; Rojec *et al.* 2019). Almost all archaeal histones lack tails and PTMs have yet to be reported. Many archaea do, however, encode multiple histone paralogs (Adam *et al.* 2017; Henneman *et al.* 2018) that can flexibly homo- and heterodimerize and – in principle – generate chromatin states of considerable combinatorial complexity.

**Figure 1.**
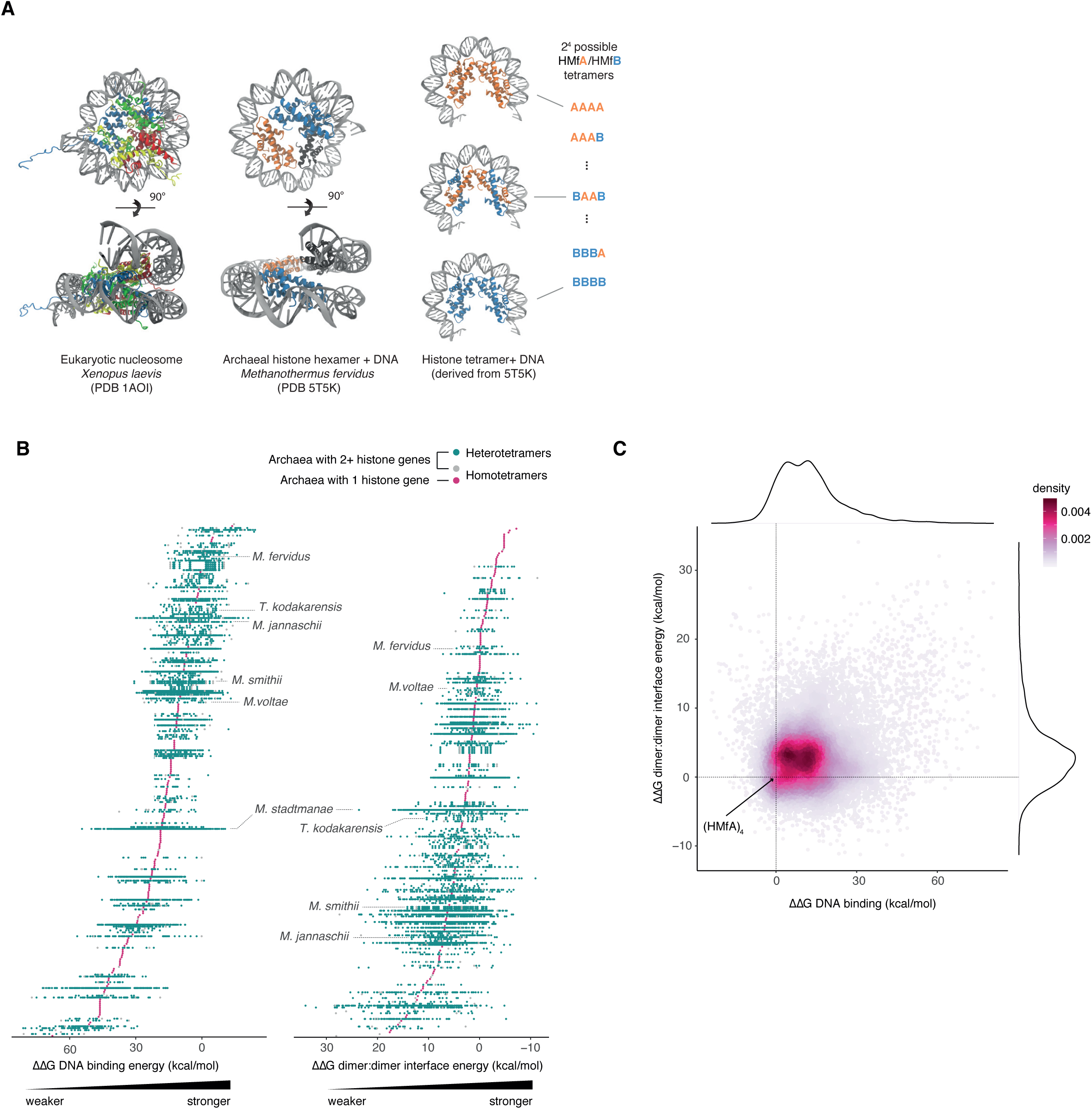
Structural diversity of archaeal histone tetramers. (**A**) Crystal structures of the octameric eukaryotic nucleosome (PDB: 1AOI), the hexameric archaeal nucleosome (PDB 5T5K), and the same structure with one dimer removed to yield the tetrameric complex, alongside a schematic showing the different combinations of histones in homo- and heterotetrameric models built for two histones (e.g. *M. fervidus* HMfA and HMfB). (**B**) DNA binding strength and tetramerization strength (dimer:dimer interface energy) for every possible tetrameric histone complex within each species of archaea in our sample. Each point, grouped by species, represents an individual complex. Species are ordered by mean interaction energy across tetramers. Species labels are provided in Figure S1. (**C**) Relationship between DNA binding and tetramerization strength for each tetrameric model. Most complexes have slightly weaker tetramerization strength and DNA binding than HMfA. ΔΔG is given relative to the HMfA homotetramer for all plots.

Prior studies in a handful of model species found that archaeal histone paralogs can differ in their expression through the growth cycle, DNA binding affinity, and oligomerization potential, and described specific effects on growth and transcription when different paralogs from the same archaeon were deleted (Sandman *et al.* 1994; Bailey *et al.* 2002; Heinicke *et al.* 2004; Cubonova *et al.* 2012; Henneman *et al.* 2018). Yet how the properties of different histone paralogs combine within a single cell to generate dynamic, responsive chromatin states and whether archaeal histone paralogs play conserved roles akin to eukaryotic histone variants remains unknown.

Here, we shed light on the evolution of archaeal histone paralogs and their capacity to generate diverse chromatin states through multimeric assembly. Combining *in silico* fast mutational scanning with molecular dynamics simulations and evolutionary analysis, we show that histone paralogs can generate substantial diversity when it comes to key structural properties of the histone-DNA complex. Using *Methanosphaera stadtmanae –* which encodes an unusually large number of histone paralogs (7) – as a case study, we show that chromatin state space in this multi-histone system is large but dense and can be traversed smoothly by altering the dosage of individual paralogs. At the same time, we highlight the potential for more radical change, describing the widespread existence of *capstones* – histones that, when incorporated, are predicted to prevent further oligomer extension. Importantly, we show that capstones (and other paralogs) in the Methanobacteriales are related by vertical descent, providing evidence for long-term maintenance of functionally distinct paralogs akin to eukaryotic histone variants. Finally, we trace divergent paralog properties to individual amino acid residues and show that paralog diversification has been driven by substitutions at structurally sensitive sites. We propose that paralog exchange might constitute a major mechanism of chromatin state change in archaea; a mechanism that was complemented – and arguably superseded – in eukaryotes by the proliferation of post-translational modifications. Our results suggests that the last common ancestor of eukaryotes, which emerged from within the Archaea (Eme *et al.* 2017; Williams *et al.* 2020), might have inherited histone-based chromatin of considerable combinatorial complexity from its archaeal ancestor, with implications for the contribution of histones to the establishment of eukaryotes (Brunk and Martin 2019).

## RESULTS

### Heteromeric histone-DNA complexes exhibit large differences in DNA binding affinity and stability across archaea

Our current knowledge of functional differences amongst archaeal histone paralogs is limited, especially for archaea with more than two histone genes, where functional diversity might be greatest. Many of these archaea remain genetically inaccessible and/or difficult to culture, pre-empting detailed experimental characterization. This includes archaea from the Asgard clade, the closest known relatives of eukaryotes (Eme *et al.* 2017; Williams *et al.* 2020). To shed light on the functional diversity of histone paralogs in archaea, we therefore combined structural modelling approaches with evolutionary analysis.

First, using the hexameric crystal structure of HMfB from *M. fervidus* as a template, we built models of tetrameric histone complexes bound to DNA for 282 diverse archaea (see Methods). Tetramers constitute the minimal oligomeric unit capable of wrapping DNA and have been widely observed in a range of archaea *in vivo* (Marc *et al.* 2002; Reeve *et al.* 2004; Mattiroli *et al.* 2017; Rojec *et al.* 2019). For archaea with more than one histone gene, we modelled all possible tetrameric combinations (*n*^4^, where *n* is the number of paralogs; Figure 1A), only excluding histones with large insertions, deletions or terminal extensions (tails) and those with deletions in the core histone fold (see Methods). This resulted in 349 homo-oligomeric and 15905 hetero-oligomeric complexes in total. We then considered Gibbs free energy changes (ΔG) at the DNA-protein interface (a measure of DNA binding affinity) and at the interface between the two histone dimers (a measure of tetramer stability, see Methods). Across our diverse sample of archaea, we observe substantial apparent variability in DNA binding affinity and tetramer stability (Figure 1B,C; Figure S1; Table S1). *Effective* differences between species might, however, be less pronounced than they appear. In fact, while we model tetrameric complexes under standardized conditions (see Methods), archaea differ widely with regard to growth temperature, pH, the concentration of organic and inorganic solutes, and other factors that can influence protein-protein and protein-DNA interactions *in vivo*. As attempts to systematically control for such potential confounders are plagued by incomplete information, we focus first on comparisons *within* species, where different heteromeric complexes can be compared more fairly. In particular, we consider *Methanosphaera stadtmanae*, a mesophilic methanogen that inhabits the human gut, as a case study.

### Methanosphaera stadtmanae *as a case study for combinatorially complex chromatin*

*M. stadtmanae* DSM3091 encodes seven non-identical histone genes, located around the chromosome as apparent single-gene operons (Figure S2). The 7^4^ (=2401) tetrameric histone-DNA complexes we built from these histones span the largest DNA affinity range (ΔΔG of - 10.47 to 54.39 kcal/mol relative to HMfA) and fourth largest tetramer stability range (ΔΔG of −9.33 to 23.63 kcal/mol) in our sample (Figure 1B,C; Figure 2A), providing an excellent model system to interrogate the capacity of an individual archaeal cell to generate different chromatin states by altering the composition of histone-DNA complexes via paralog exchange.

**Figure 2.**
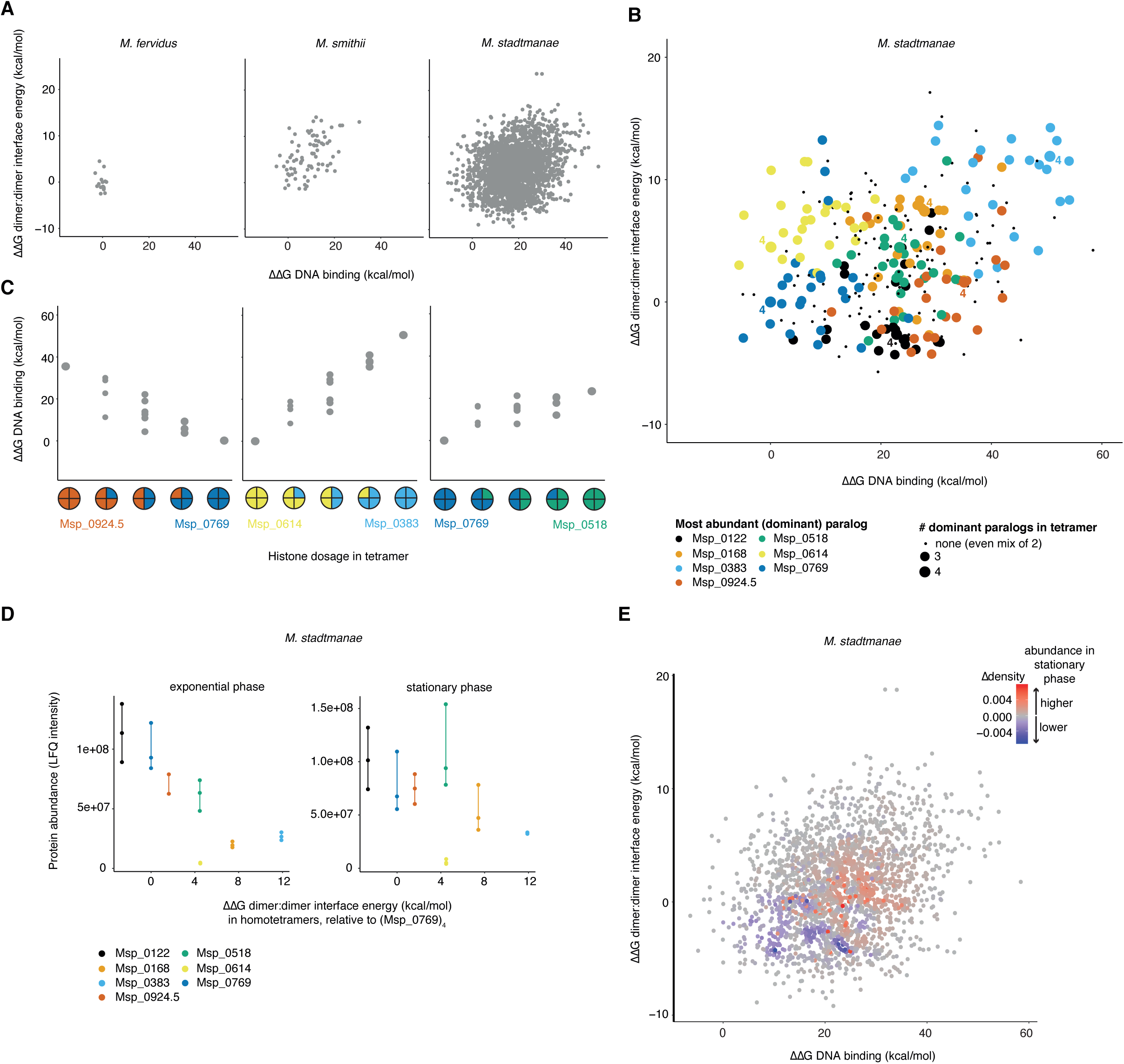
Structural diversity of *Methanosphaera stadtmanae* tetrameric histone-DNA complexes. (**A**) Chromatin state space defined by DNA binding and tetramerization strength for models built using fast mutational scanning for *M. fervidus* (2 histone paralogs), *M. smithii* (3 histone paralogs) and *M. stadtmanae* (7 histone paralogs). ΔΔG is given relative to the HMfA homotetramer. (**B**) Chromatin state space defined by DNA binding and tetramerization strength for *M. stadtmanae* histone complexes containing ≤ 2 histone paralogs. Points are coloured by the dominant paralog in the complex (3 or 4 out of 4 monomers in the tetramer). Homotetramers are labelled (“4”). (**C**) Examples of DNA binding strength varying gradually with paralog dosage. (**D**) Tetramerisation strength of *M. stadtmanae* homotetramers compared to empirically determined paralog abundance in exponential and stationary phase. (**E**) Relative change in the abundance of different tetrameric complex in stationary versus exponential phase, as predicted by sampling 100,000 tetrameric complexes based on relative protein abundance (mean LFQ intensity) in exponential and stationary phase. Increased abundance of complexes in stationary phase is shown in red, decreased abundance in blue. ΔΔG is given relative to Msp_0769 for panels B, C, D and E.

We find that tetrameric combinations are not randomly distributed across this state space but occupy partially distinct areas based on which paralog dominates the complex (Figure 2B). Homotetramers are found towards the edges of this space while the intervening space is densely populated (Figure 2A,B). Complexes that are intermediate in terms of paralog dosage, tend to have intermediate properties, enabling smooth transitions in chromatin state space, from one extreme to another (Figure 2B,C). Paralogs in this system therefore provide the capacity for graded control of chromatin state through changes in relative paralog dosage, as well as for more radical transitions (see below).

### *In vivo expression dynamics of histone paralogs in* Methanosphaera stadtmanae

Is the capacity for graded control of chromatin state used dynamically *in vivo*? And what areas of chromatin state space are actually explored? To begin to address the latter question, we quantified the relative abundance of histone paralogs in exponential and stationary phase *M. stadtmanae* cells using label-free mass spectrometry and RT-qPCR (see Methods, Figure S3). Protein abundance varies over a 27-fold range between paralogs but expression levels of individual paralogs are well correlated in exponential and stationary phase (Figure 2D, Figure S3). Intriguingly, relative paralog abundance in exponential phase exhibits a strong correlation with tetramer stability: paralogs that are inferred to form more stable homotetramers are more abundant (rho=-0.82, P=0.034; Figure 2D). This is also the case (based on previously determined relative transcript abundances) in *Methanobrevibacter smithii*, another member of the order Methanobacteriales, but not in the hyperthermophiles *M. fervidus, T. kodakarensis*, and *Thermococcus onnurineus* (Figure S4).

To mimic the relative abundance of different complexes in the cell and better approximate actual vis-à-vis theoretical chromatin state space *in vivo*, we generated 100,000 tetrameric complexes *in silico*, with individual histones recruited into each complex at random based on their relative abundance at the protein level. Assuming that histones dimerize randomly, we find that the centre of gravity in chromatin state space shifts towards complexes that are, on average, less stable, exhibit lower DNA binding affinity (Figure 2E, Figure S5), and therefore likely give rise to fewer stable higher-order oligomers. This shift is driven by the upregulation of two histones, Msp_0168 and Msp_0518 (Figure 2D, Figure S3), which we infer exhibit relatively low DNA binding affinity and tetramer stability as homotetramers. Thus, we predict that stationary phase should – other things equal – be characterized by more open histone-based chromatin. This again contrasts with prior observations in *M. fervidus*, where expression of HMfB – capable of greater DNA compaction - increases in stationary phase relative to HMfA, the second paralog in *M. fervidus* (Sandman *et al.* 1994). We return to this difference below.

### *Molecular dynamics simulations of* M. stadtmanae *homotetramers*

To gain more detailed insights into the extremes of *M. stadtmanae* chromatin state space, we carried out extensive molecular dynamics simulations on all its homotetrameric histone-DNA complexes. DNA binding affinities inferred from these simulations correlate well with results obtained from fast mutational scanning (rho=0.96, P<0.001, Figure S6), providing further validation that the fast mutational scanning approach captures salient properties of the histone-DNA complex. But the simulations also reveal additional details. Most notably, we find that Msp_0383, the paralog with the lowest predicted DNA binding affinity and tetramer stability, exhibits much more extreme spatial displacement from the starting point of the crystal structure than the other histones (Figure 3A). Further analysis of trajectories over the 100 ns simulation revealed that the Msp_0383 homotetramer displays an unstable dimer:dimer interface – unlike the other homomeric complexes, which reach an approximate equilibrium after <20 ns. While the two Msp_0383 dimers remain individually bound to DNA, they are refractory to tetramerization (Figure 3A, File S1-2). Thus, our modelling predicts that Msp_0383 assembles into histone-DNA complexes that are structurally distinct from classic tetrameric complexes observed for *M. fervidus*, other model archaea, and the remaining *M. stadtmanae* paralogs.

**Figure 3.**
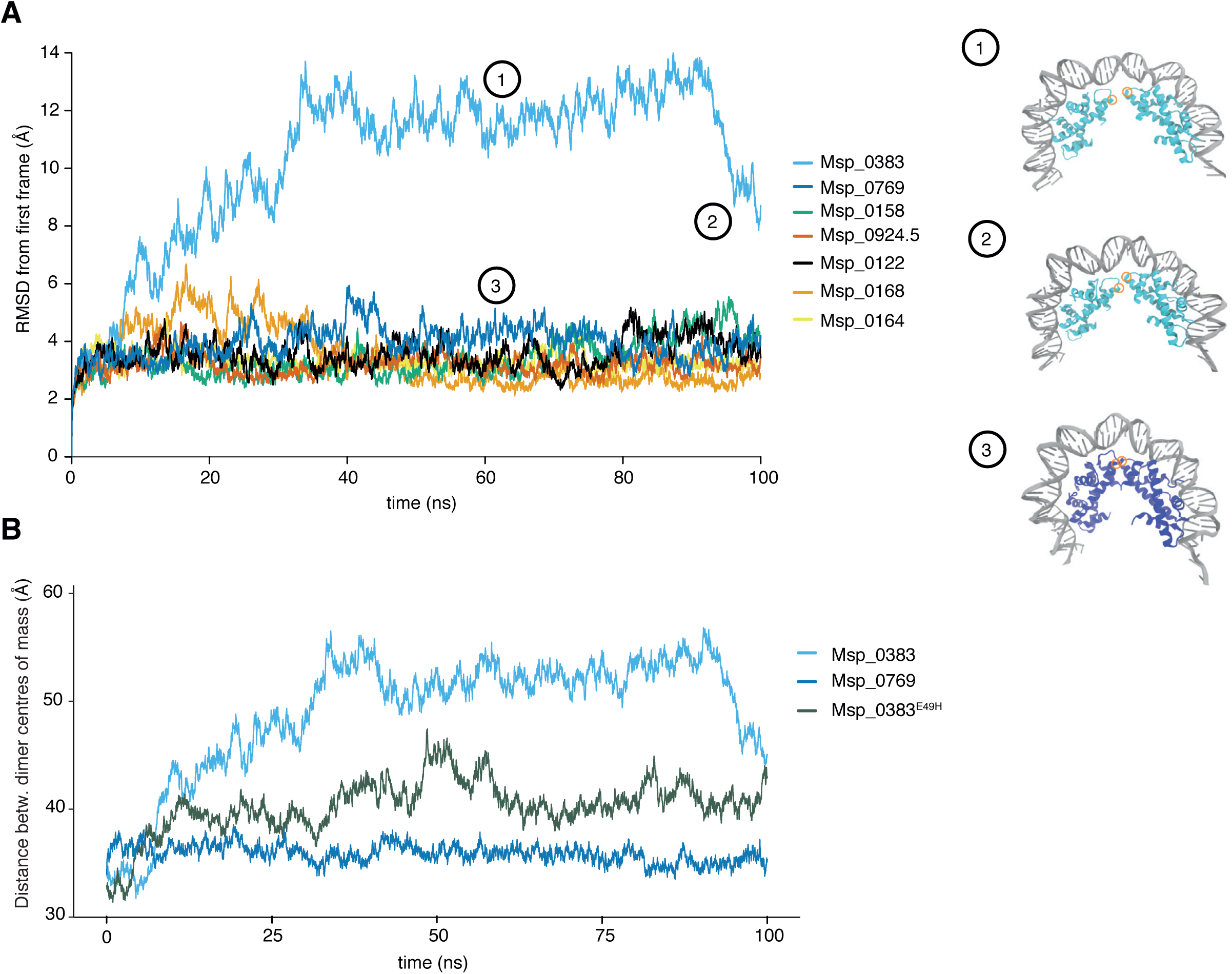
Molecular dynamics simulations of homotetrameric histone models from *M. stadtmanae*. (**A**) Root mean square deviation (RMSD) over the course of the simulation, beginning from the crystal structure into which *M. stadtmanae* histones have been substituted (see Methods). Example structures at different frames are shown for Msp_0383 and Msp_0769 with the position of residue 49 (residue 50 in Msp_0383) highlighted in orange. (**B**) Distance between the centre of mass of dimers in homotetrameric models of Msp_0383, Msp_0769 and the mutant Msp_0383^E49H^.

Msp_0383 has a negatively charged glutamic acid (E) at position 49 whereas the other paralogs (and most histones across archaea) have a positively charged histidine (H, Figure 3A; note that, throughout the manuscript, we number residues based on positional orthology to HMfB; the raw residue number in *M. stadtmanae* is 50). Residue 49 is close to the interface between dimers and mutations at this site were previously shown to impact tetramer formation of HMfB *in vitro* (Marc *et al.* 2002). To test whether amino acid identity at this site is sufficient to account for the repulsive effects observed, we *in silico*-substituted E for H in all histones of the tetramer and subjected the resulting complex to the same simulation protocol. We find that this substitution alone is enough to significantly reduce the distance between dimers, with Msp_0383^E49H^ exhibiting dynamics that are intermediate between Msp_0383 and the other paralogs (Figure 3B). These results suggest that Msp_0383 functions as a capstone, preventing tetramerization and, when tagged onto an existing complex, further oligomerization.

Such potential capstones are not unique to *M. stadtmanae* but also present in other members of the Methanobacteriales, as demonstrated by comparative molecular dynamics simulations of *M. smithii* homotetramers, which also reveal a single, lowly expressed histone (Msm_1260) associated with much-reduced tetramer stability (Figure S4,7).

### Phylogenetic analysis reveals long-term persistence of archaeal histone variants

Some eukaryotic histone variants are ancient and have persisted through multiple rounds of speciation as recognizable, distinct paralogs, often with conserved function and dedicated chaperones that can discriminate between them (Talbert and Henikoff 2010). Notably, this includes H2A.Z, which emerged at the base of eukaryotes. Other variants, like macroH2A are restricted to certain clades and therefore evolved more recently. Yet others, like H2A.X, appear polyphyletic in origin, pointing to repeated independent emergence of functionally analogous variants (Malik and Henikoff 2003; Talbert and Henikoff 2010). What is the situation for archaeal histones? Are there persistent, recognizable paralogs of ancient origin? Or is most diversification relatively recent and lineage-specific? We know that histones with predicted capstone properties are found in various archaeal genomes [see above and (Henneman *et al.* 2018)]. But did capstones have a single origin?

Phylogenetic analysis of histones across archaea is complicated by the fact that histones are short (<70 amino acids) and timescales are large, leading to poorly supported nodes in a global phylogeny of archaeal histones (see Methods, File S3-8). We therefore focused our analysis on the Methanobacteriales, which include Methanosphaera, Methanobrevibacter, and Methanobacterium *spp.* as well as *M. fervidus* (Figure 4A). Alongside abundant lineage-specific duplication events (shaded taxon labels in Figure 4B,C), we find several cases of longer-term paralog maintenance, indicated by the existence of multiple groups of sequences that each recapitulate the species phylogeny and further supported by conserved synteny. For example, branching patterns and synteny of histones in Methanobacterium strongly suggest two ancient gene duplication events that preceded the divergence of this genus (Figure 4B). Importantly, synteny analysis also reveals maintenance of paralogs *between* Methanobrevibacter and Methanobacterium (groups 1 and 3 in Figure 4B,C), indicating that these originated from even more ancient duplications, dating back to the ancestor of these two genera. As synteny breaks down further, making confident assignments becomes harder. Closer inspection of local gene neighbourhoods, however, suggests that there might be even deeper conservation of recognizable paralogs all the way out to *M. fervidus*, where HMfB (HMfA) is flanked upstream (downstream) by *trpS* (*radB*), whose relative position is conserved in Methanobrevibacter and Methanobacterium *spp*. (Figure 4B,C). Finally, regarding capstones, we find evidence for shared vertical descent of the *M. stadtmanae* and *M. smithii* capstones (Figure 5), providing evidence for long-term maintenance of distinct histone functionalities in archaea.

**Figure 4.**
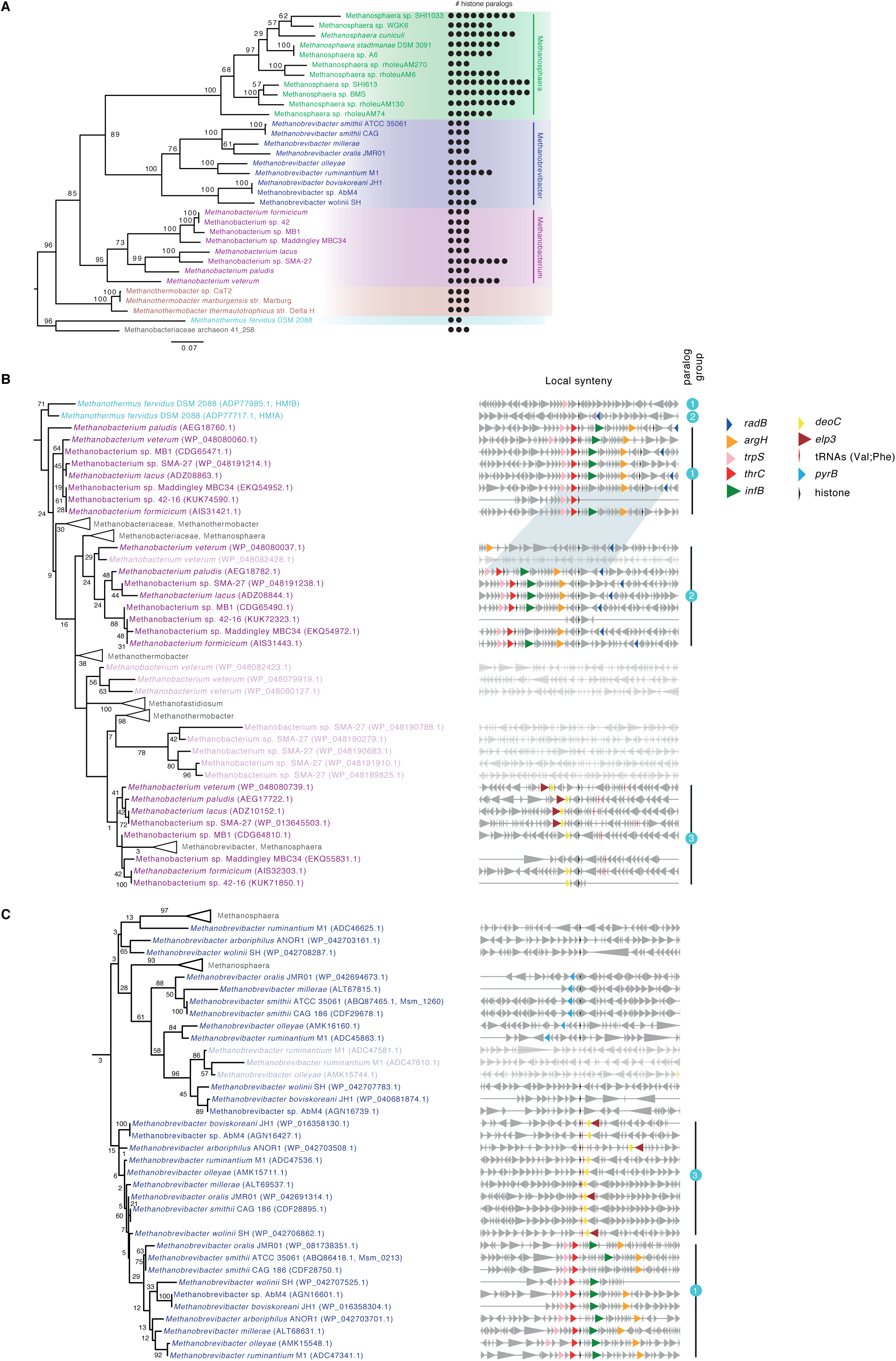
Phylogenetic analysis of histone paralogs in the Methanobacteriales. (**A**) Maximum likelihood reference phylogeny (species tree) of the order Methanobacteriales, using IF-2a as a representative, vertically inherited gene. Bootstrap values are shown as a percentage out of 200 non-parametric bootstraps. The tree is rooted with *M. fervidus* as the outgroup. The number of histone paralogs in a given genome is mapped on the right-hand side. (**B**) Examples of recent duplications (light-shaded taxon labels) and long-term maintenance of paralogs in the genus Methanobacterium and (**C**) Methanobrevibacter, as supported by tree topology and conserved synteny. Note the clustering of proteins according to shared synteny rather than species phylogeny. Examples of paralog groups (1-3) are highlighted. Shared synteny across the Methanobacterium/Methanobrevibacter divide supports paralogous relationships (group 1 & 3). Even deeper paralogy is suggested by the fact that local spatial association of *trpS* (group 1) and *radB* (group 2) with histones extend to *M. fervidus.* Longer gaps in the synteny blocks, evident for individual genomes, are the result of incomplete genome annotations. Genes are automatically color-coded based on similarities in functional annotation (see Methods). The trees are rooted with reference to a wider phylogeny of archaeal histones (see Methods) and bootstrap values shown as a percentage out of 500 non-parametric bootstraps. In both (B) and (C) some sequences from other Methanobacteriales have been collapsed for clarity. The scale bar represents the average number of substitutions per site. Full trees and the underlying alignments are provided as File S3-8.

**Figure 5.**
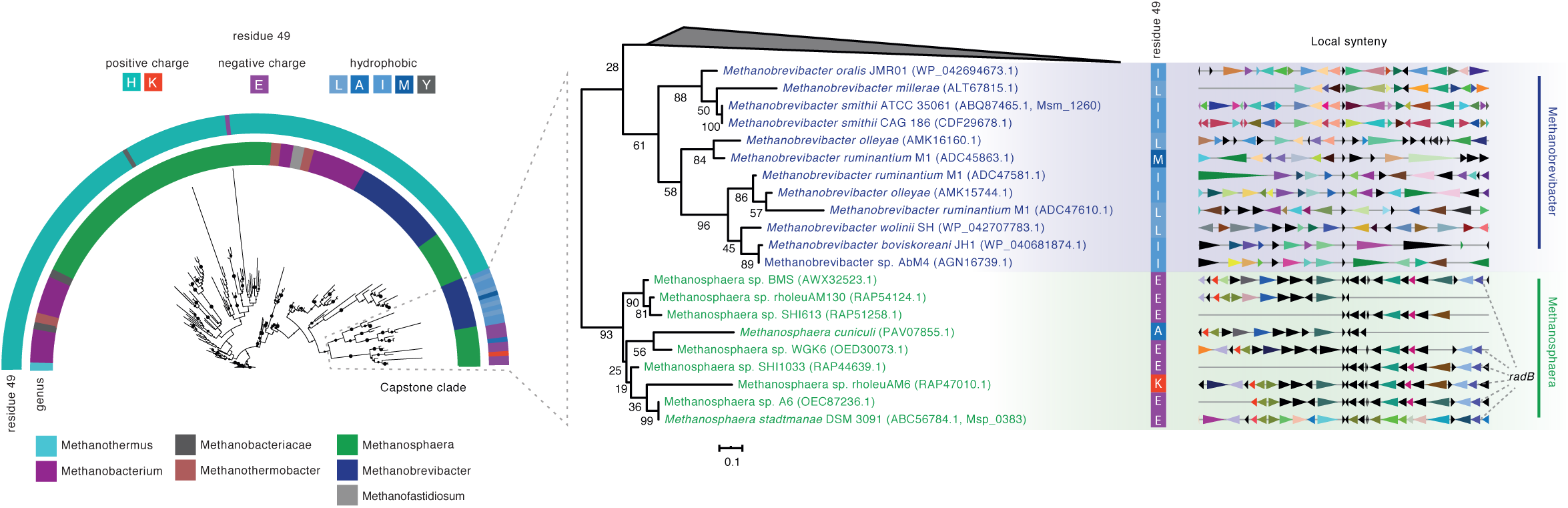
Phylogenetic analysis of capstone histones. A maximum likelihood tree including all Methanobacteriales genomes (File S8) is displayed on the left, along with information on amino acid identity at residue 49. Histone paralogs with capstone properties (negatively charged or hydrophobic amino acids) in Methanosphaera and Methanobrevibacter *spp.* cluster to the exclusion of other histones found in these species. Local synteny in the vicinity of histone paralogs is shown on the right. Genes are automatically color-coded based on similarities in functional annotation. The *radB* gene is highlighted to allow cross-referencing with Figure 4B,C. Bootstrap values are shown as a percentage out of 500 non-parametric bootstraps. The scale bar represents the average number of substitutions per site.

To put the time-scale of paralog origin into context, we note that the lineages leading to *M. stadtmanae* and *M. smithii* split an estimated ∼1.3Gya, while the wider Methanobacteriales are thought to have emerged as a clade approximately ∼1.6Gya (Wolfe and Fournier 2018). At least some archaeal histone variants have therefore been maintained for hundreds of millions of years of evolution, rendering them comparable in age to the oldest known eukaryotic histone variants, which date back to the last common ancestor of eukaryotes roughly 1.2-2Gya (Eme *et al.* 2014; Betts *et al.* 2018).

### Single amino acid changes underpin functional differences between paralogs

The case of *M. stadtmanae* Msp_0383 illustrates that substitutions of individual amino acids can have strong effects on histone properties and, ultimately, chromatin state. This is also true in eukaryotes (Maze *et al.* 2014; Henikoff and Smith 2015; Nacev *et al.* 2019). H3.3 and H3.1, for example, differ in only four amino acids (three of which are located in the histone fold domain), but are recognized by different chaperones, deposited at defined locations along the genome, and make distinct, non-redundant contributions to genome function, notably during gametogenesis (Talbert and Henikoff 2010; Filipescu *et al.* 2014; Wollmann *et al.* 2017).

To understand how specific amino acid changes underpin the functional diversification of archaeal histone paralogs, we integrated structural modelling and evolutionary analysis. First, we used the FoldX forcefield (see Methods) to *in silico-*mutate each amino acid in the model histone HMfB from *M. fervidus* to every other possible amino acid to identify regions and individual sites particularly sensitive to change. We then compared these predicted effects to previous *in vitro* work on HMfB, which had identified residues that, when mutated, affect DNA binding, the direction of DNA supercoiling, rigidity of the histone-DNA complex, thermostabilisation, oligomer formation, and the ability of the histone to accumulate in *E. coli*, a proxy for folding stability (Soares *et al.* 2000; Marc *et al.* 2002; Soares *et al.* 2003; Higashibata *et al.* 2003; Mattiroli *et al.* 2017). We find that predicted and observed effects are highly concordant (Figure 6A,B; Table S2). For example, our fast mutational scanning identifies the four residues (46, 49, 59, 62; Figure 6D) previously highlighted as critical for stable tetramerization (Marc *et al.* 2002; Henneman *et al.* 2018) and we correctly predict which mutations had led to increased (stronger DNA binding) or decreased (weaker DNA binding) mobility in a previous gel shift assays (Figure 6B). This high degree of congruence provides additional validation for our modelling approach. It also increases our confidence in predictions of structural sensitivity for residues that have not been experimentally interrogated. For example, residues 21 and 50, for which no experimental data is available, show large deviations in DNA binding affinity and tetramerization strength, respectively, when mutated (Figure 6D).

**Figure 6.**
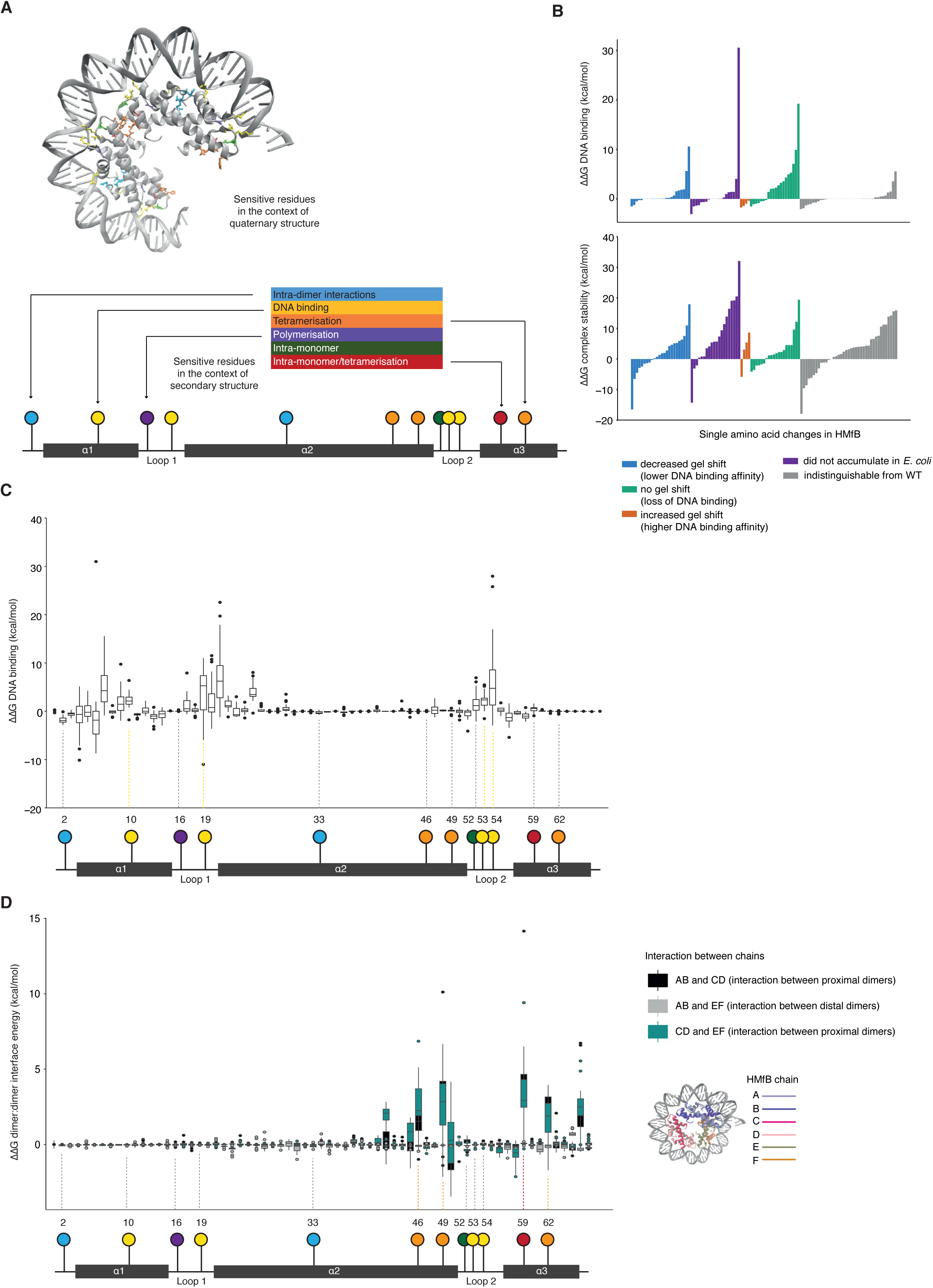
Modelling the impact of single amino acid mutations on the (HMfB)_6_-DNA complex. (**A**) Residues where mutations are known from previous experimental work (see references in Introduction) to affect monomer:monomer interactions, DNA binding, tetramerisation, polymerisation and intra-monomer interactions are highlighted on the quaternary and secondary structure. (**B**) FoldX-calculated changes in DNA binding affinity (top) and stability (bottom) for HMfB single amino acid mutants previously characterised qualitatively in gel shift experiments (Soares *et al*. 2000). Individual mutations are listed in Table S2. (**C**) DNA binding and (**D**) tetramerization strength for all possible single amino acid mutations of HMfB. The location of residues with previously known function is shown on the secondary structure beneath. For (D), the resulting interaction energy between each dimer pair in the hexamer was calculated and the location of dimer pairs in the hexamer is shown. ΔΔG is quoted relative to the wild-type HMfB structure for all plots.

Next, we asked how this comprehensive landscape of possible effects compares with substitutions that actually occurred during the evolution of archaea. Do structurally sensitive sites remain largely conserved across paralogs? Or are changes at key sites, like those we observe for Msp_0383, relatively commonplace? To answer this question in a pan-archaeal manner, we took a non-phylogenetic approach. We aligned the 506 archaeal histone proteins in our sample (see Methods) and then split them into two groups, depending on whether they come from a genome that encodes only a single histone gene or from a genome that encodes two or more paralogs. Our objective here was to identify residues along the histone fold that have become more diverse in multi-paralog systems, where relaxed constraint or positive selection could drive diversification following duplication. Figure 7 shows the amino acid diversity ratio *H*_*M*_/*H*_*s*_ for each residue, where *H*_*M*_ and *H*_*s*_ are Shannon diversity indices calculated for a given residue (column in the alignment) across multi-histone and single-histone genes, respectively (see Methods). The average Shannon ratio will be affected by phylogenetic sampling, the number of histones in each group, and other factors, and is therefore relatively uninformative. What is informative, however, are deviations from this average at specific residues.

**Figure 7.**
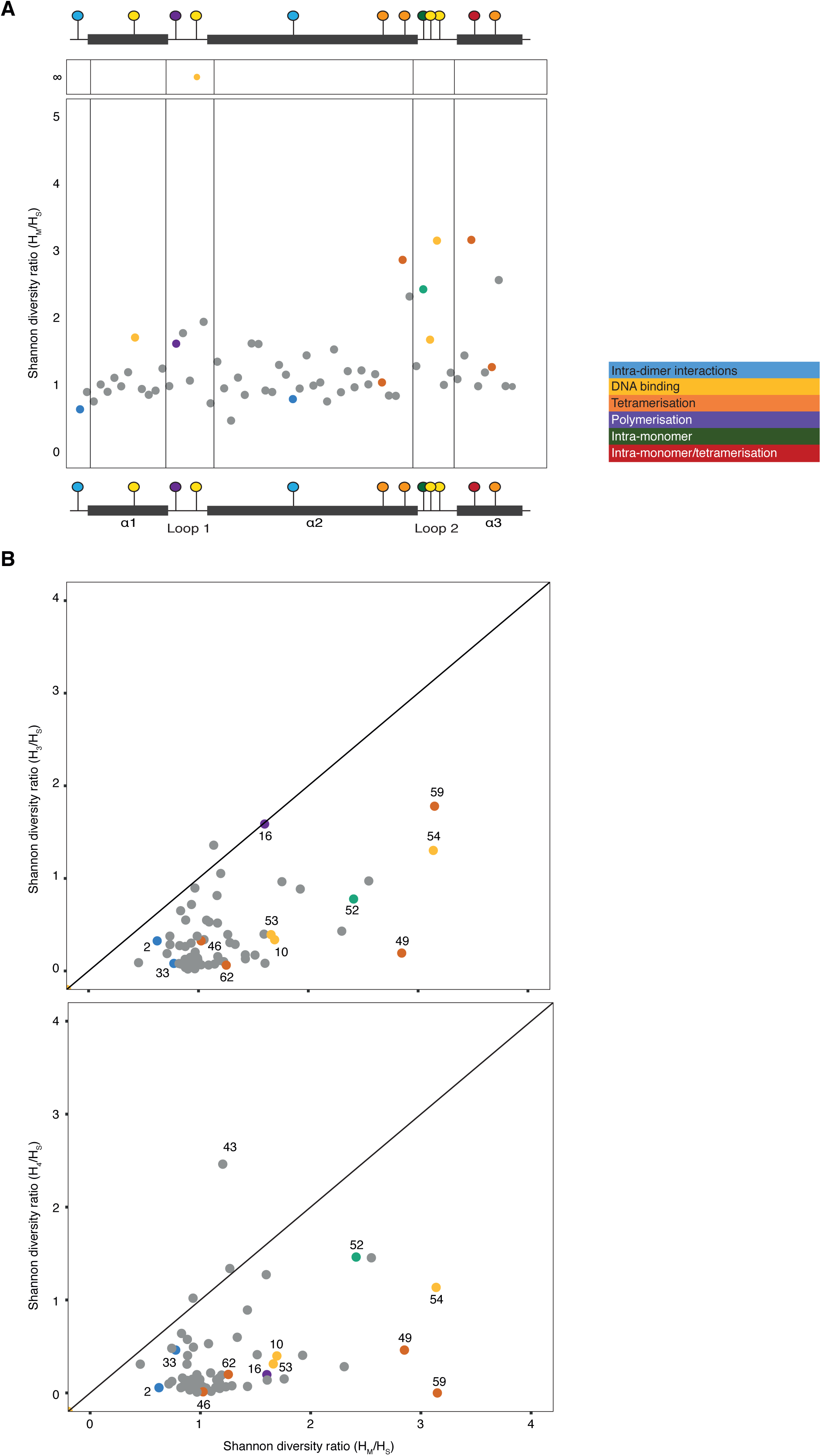
Comparative analysis of sequence diversity in archaeal and eukaryotic histones. **(A)** Shannon diversity ratio (*H*_*M*_*/H*_*S*_) at each position in the core histone fold domain. Residues are coloured by key function from previous mutational studies (see Figure 6). *H*_*S*_ for residue 19 is 0, so the Shannon ratio is undefined. (**B**) Shannon diversity ratios for H3 (*H*_*3*_*/H*_*S*_, top panel) and H4 (*H*_*4*_*/H*_*S*_, bottom panel) compared to *H*_*M*_*/H*_*S*_. Residues of particular interest are numbered.

Strikingly, diversification in species with multiple paralogs is strongly associated with structurally critical residues (Figure 7A). This includes the capstone residue 49, but also several residues that make large contributions to DNA binding (10, 19, 53, 54; Figure 6C,D, Figure 7A), concentrated in the loop regions of the histone fold, and loop 2 in particular. Perhaps the most egregious example is residue 19 in loop 1, which is perfectly conserved as an arginine in single-histone archaea (*H*_*s*_=0) but accommodates eight different amino acids across the multi-histone archaea in our sample. This suggests a significant change in the evolutionary regime at this site once more than two histones are present in the system. Given the strong deviation from the baseline diversity ratio, we think that positive selection is likely implicated in the diversification process rather than relaxed constraint alone.

Although we do not explore this extensively here, we note that residue-level diversification has some phylogenetic structure. Whereas some residues, including residue 49, carry diverse amino acids in genomes from multiple independent lineages, others exhibit a narrower phyletic pattern (Figure S8). Notably, this is the case for residue 54, which forms a conserved interaction with residue 19 (Mattiroli *et al.* 2017). Diversification at this residue is confined almost entirely to the Asgard clade and excluding this clade from the analysis dramatically reduces diversity at residue 54 (Figure S9).

### Diversification of eukaryotic versus archaeal histone folds

The residues involved in archaeal histone tetramerisation are also important for interactions at the interface of two H3 molecules from neighbouring H3:H4 dimers (Luger *et al.* 1997; Postberg *et al.* 2010). How, then, do archaeal histone paralogs compare to eukaryotic histone variants? Did diversification of the core histone fold follow a similar path? To address this question, we first added eukaryotic H3 and H4 sequences to our pre-existing alignment of archaeal histones (see Methods, Table S3). We then calculated Shannon diversity indices for H3 (*H*_*3*_) and H4 (*H*_*4*_) proteins found across eukaryotes and compared *H*_*4*_*/H*_*s*_ and *H*_*3*_*/H*_*s*_ to *H*_*M*_*/H*_*s*_. We find that diversification dynamics across the histone fold follow a similar pattern in multi-histone genes and H3 (rho=0.40, P=0.0081) and to a lesser extent also H4 (rho=0.24, P=0.063). Residues 2 and 33, which are involved in intra-monomer interactions, are not diverse in either H3, H4 or archaeal histones. Substitutions at these positions may prevent the formation of the tertiary histone fold structure and are therefore selected against. Conversely, residues around the loop 2 region in particular experience accelerated diversification in both H3 and multi-histone archaea relative to single-histone archaea. These similarities notwithstanding, several residues show conspicuous diversification in multi-archaeal histones but not H3/4, and vice versa. This includes residues 49 (high *H*_*M*_*/H*_*s*_, low *H*_*3/4*_*/H*_*s*_), 59 (high *H*_*M*_*/H*_*s*_, low *H*_*4*_*/H*_*s*_), and 43 (low *H*_*M*_*/H*_*s*_, high *H*_*4*_*/H*_*s*_). In addition, even residues with high diversity ratios in both eukaryotes and archaea only partially explore the same part of sequence space and tend to evolve towards different sets of amino acids (Figure S9). Our results therefore suggest that histone variants from archaea and eukaryotes independently focused their exploration of structural-functional space on structurally sensitive sites in the loop 2 region but also highlight significant lineage-specific constraints on histone evolvability.

## DISCUSSION

Prior observations – from variable expression along the growth cycle to differential phenotypic effects upon deletion (Sandman *et al.* 1994; Heinicke *et al.* 2004; Cubonova *et al.* 2012) – pointed to functional diversity of archaeal histone paralogs. The observations we report here not only reinforce this notion but demonstrate that some histone paralogs in archaea have been maintained as distinct functional units over long evolutionary time-scales, akin to eukaryotic histone variants. Our modeling results suggest that paralogs, by exploiting the combinatorial opportunities of histone oligomerization, can generate diverse chromatin states at the level of individual histone-DNA complexes and enable both subtle, graded dosage-driven transitions and more radical changes such as those associated with the expression of capstones.

We explored one of these transitions in *M. stadtmanae*, where the relative expression of different histone paralogs changes in stationary versus exponential phase. Based on our structural modelling and empirical protein abundance data, we predict that stationary phase in *M. stadtmanae* (as well as *M. smithii*) should be characterized by a larger fraction of less stable histone-DNA complexes. This is, arguably, unexpected given opposite trends inferred for *M. fervidus* and other hyperthermophiles and the general notion that stationary phase is associated with greater chromatin compaction. Experimental data will ultimately be required to determine whether this inferred difference is genuine or not. However, the discrepancy serves as a timely reminder to highlight the limitations of our modelling approach, which does not consider absolute histone titres, changes in intracellular conditions (e.g. in terms of solutes), and expression of other abundant architectural proteins (e.g. Alba) that will co-determine higher-order chromatin states. In this regard, our results should be considered a valuable starting point and incentive for further exploration rather than the final word, hewn in stone, on comparative chromatin complexity in archaea.

Along similar lines, we note that we only examined a small branch of the archaeal tree in depth, did not consider archaeal histone with tails or large indels (Friedrich-Jahn *et al.* 2009; Henneman *et al.* 2018) and did not explore interactions and combinatorial complexity beyond the tetramer level. Our estimates of archaeal capacity to generate different chromatin states is therefore likely conservative. In particular, tetramer models do not allow us to consider stacking interactions, which affect oligomerization propensity (Mattiroli *et al.* 2017; Henneman *et al.* 2018). Substantial additional complexity might further emerge from the consideration of N-terminal tails, which are present in some Heimdallarchaea (Mattiroli *et al.* 2017; Henneman *et al.* 2018), the closest known relatives of eukaryotes (Williams *et al.* 2020). Studying these archaea and their tails will be particularly important to understand what – in the context of histone-based chromatin – constitutes eukaryotic innovation, elaboration or shared archaeal heritage.

Based on our current knowledge, we speculate that paralog-mediated structural change might play an outsize role in archaea compared to eukaryotes, where post-translational modifications and interactions with other proteins are heavily involved in altering chromatin state in response to upstream signals. One of the key eukaryotic innovations might have been a switch from predominantly paralog-based generation of different chromatin states to using an octameric nucleosome as a platform for integrating epigenetic information. This innovation might also have enabled another: local specification. In eukaryotes, divergent regulatory states can be encoded along the same chromosome via targeted deposition of paralogs and histone marks by enzymes and chaperones that can interact with specific histones, DNA sequences, and/or other constituents of chromatin. At present, we have no evidence that the capacity for such local control exists in archaea. Current data only support a global, genome-wide role in re-shaping chromatin state. It will be interesting in the future to determine whether complexes of different composition are indeed randomly distributed or show non-random patterns along archaeal chromosomes in a manner anticipating eukaryotic chromatin. To this end, we need to develop a better understanding of archaeal histone variants in physiological context. The specific functional roles of archaeal variants in the context of genome function remain entirely unknown, a glaring gap that can only be plugged by *in vivo* experiments. Our study provides ample incentive for further research to establish how archaeal paralogs are regulated, how they interact with other DNA-binding proteins to determine global and perhaps local chromatin states, and how paralogs contribute to adaptive responses in physiological context.

## METHODS

### Alignment of histones

A previously compiled set of archaeal histone proteins (Adam *et al.* 2017) was filtered to only include proteins between 60-80 amino acids in length with a single histone fold (Figure S10). For reference, HMfB is 69 amino acids long. This filtered set of histones from 282 species of archaea (139 with more than one histone, 143 with one histone) was aligned using MAFFT-linsi (-localpair-maxiterate 1000) (Kazutaka Katoh 2013). Eukaryotic H3 and H4 protein sequences were downloaded from InterPro (matching folds IPR007125 and IPR032454) (Mitchell *et al.* 2019) and filtered for length; 95-110 amino acids for H4, 130-145 amino acids for H3 (Figure S10). Sequences were further filtered to randomly remove redundant entries (i.e. sequences 100% identical to another entry) and then added to the archaeal histone alignment using MAFFT-linsi (-localpair -maxiterate 1000 -seed). Positions where more than 5% of sequences had a gap were removed from further analysis.

### HMfB single mutants

We used the BuildModel command in FoldX (Schymkowitz *et al.* 2005) to introduce all possible single amino acid changes into the HMfB hexamer (PDB structure 5T5K). All six histone monomers in the structure were mutated simultaneously. FoldX refines structures by minimising the energy of mutated side-chain residues and neighbouring residues according to its empirically derived forcefield. The positions of non-adjacent residues and all peptide backbone atoms remain fixed. Although molecular dynamics simulations are more rigorous to determine accurate binding affinities and allow us to sample the dynamics of the complex (see below), FoldX allows us to sample, at high-throughput, changes in energy associated with mutations at individual positions in the protein. We therefore refer to this approach as a fast mutational scanning technique. FoldX was used at the default temperature setting of 298 K. We calculated the relative change in Gibbs free energy (ΔΔG) of the system, DNA binding and tetramerization energies for each mutant using FoldX relative to the minimised HMfB hexamer structure.

The Gibbs free energy (eq 1.) is a thermodynamic quantity defined as the amount of reversible work a mechanical system can undergo. Where ΔH is the enthalpic contribution and ΔS is the entropic contribution. By calculating the sum total of inter- and intra-molecular forces, determined by the FoldX forcefield, we can calculate ΔG and predict the structural stability of the complex. By subtracting ΔG_mutant_ from ΔG_wildtype_ of HMfB we arrive at the relative change in Gibbs free energy, ΔΔG (eq 2.). The binding affinity can be determined by subtracting the energetic contribution from the DNA and histone from the complex (eq 3.). The same can be said for the histone tetramerisation energy; subtracting dimer energies from the tetramer energy will leave us with the energetic contribution of tetramerisation.

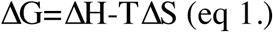

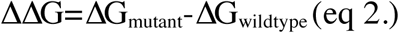

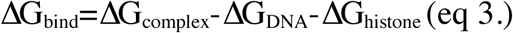

### Tetramer models of all archaeal monomeric histones

The HMfB tetramer model was built by removing one histone dimer (chains E and F) and 30bp of DNA from the 5T5K PDB structure. For each species, all possible combinations of histone monomers were modelled as a tetramer, with the following exceptions: To enable fair structural comparison, we only analysed models where no histone carried a deletion in the core histone fold (HMfB residues 2-65) and only considered histones 60-80 amino acids in length. We focused on tetramers as this allows DNA binding and tetramerisation strength to be calculated without assuming that histones assemble into longer oligomers. Histone models with deletions relative to HMfB were removed from the dataset for analysis. Substitutions were then mapped onto the HMfB tetramer using the BuildModel function of FoldX (Schymkowitz et al., 2005). Structures were energy minimised for 10000 steps combined of steepest-descent and conjugate gradient using AmberTools. Unlike FoldX, which only minimises mutated side-chain residues and their neighbours, we used an all-atom minimisation (using AMBER ff14SB) but avoided any significant refolding by applying a 2 kcal/mol/Å^2^ harmonic restraint on backbone atoms.

Binding affinity and tetramerization energies were calculated using the single trajectory MMPBSA approach (Miller *et al.* 2012). In this method we decompose ΔH in eq 1. into the gas phase energy and the free energy of solvation (eq 4.). The gas phase energy was calculated as the total of energy from the AMBER ff14SB forcefield (Maier *et al.* 2015) and the free energy of solvation was approximated using the Poisson-Boltzmann equations.

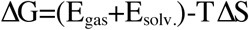

As single mutations will not significantly change the conformation of the histone, the relative change in entropy, ΔΔS, will be close to zero. For this reason, we have not included the entropic contribution to the Gibbs free energy values. ΔΔG was calculated relative to the Msp_0769 homotetramer for *M. stadtmanae* tetramer models and relative to the HMfA homotetramer in all other cases.

### Molecular dynamics simulations of Methanosphaera stadtmanae

Complexes of homotetrameric histones with DNA were parameterised using the Amber ff14SB potentials for canonical proteins using tLeap in AmberTools. Residues present in the sequence but removed in the filtering stage after alignment were manually added to the ‘full model’ homotetrameric structures generated by FoldX and the complexes were energy-minimised as above. Models were solvated with 14Å of TIP3P water and neutralised with NaCl. Energy minimisation was performed for 2000 steps using combined steepest descent and conjugate gradient methods. Following minimisation, 20 ps of classical molecular dynamics (cMD) was performed in the NVT ensemble using a Langevin thermostat (Davidchack *et al.* 2009) to regulate the temperature as we heated up from 0 to 300 K. Following the heat-up phase, we performed 100 ns of cMD in the isobaric/isothermal (NPT) ensemble using the Berendsen barostat (Berendsen *et al.* 1998) to maintain constant pressure of 1 atm during the simulation. All simulations were performed using GPU (CUDA) version 18.0.0 of PMEMD (Götz *et al.* 2012; Le Grand *et al.* 2013; Salomon-Ferrer *et al.* 2013) with long-range electrostatic forces treated with Particle-Mesh Ewald summation (Essmann *et al.* 1995).

### Phylogenetic and evolutionary analysis

To build an initial tree of archaeal histones, we queried all 282 species present in the structural analysis and available through NCBI with hmmsearch (HMMer suite, http://hmmer.org) and considered all single domain hits against Pfam model CBFD_NFYD_HMF (PF00808, Pfam v.23) that were filtered out from the initial dataset. For reproducibility purposes, the Pfam gathering threshold was used as the thresholding option of hmmsearch (--cut_ga). Sequences were first aligned using MAFFT-linsi (using blosum30) and an initial tree inferred using IQ-TREE2 (automatic substitution model estimation : LG+R6 substitution model, 1000 ultra-fast bootstraps) (Minh *et al.* 2020). We then considered the minimal subtree containing all histones from *Methanosphaera stadtmanae*. To extend the diversity of Methanosphaera histones, we downloaded additional available Methanosphaera genomes from the NCBI refseq database. All sequences were then re-aligned using MAFFT-linsi (using blosum62), and a maximum likelihood tree was build using RAxML-ng (500 non-parametric bootstrap, LG substitution model) (Kozlov *et al.* 2019). Trees were visualised using iTol (Letunic and Bork 2019) and local synteny using Genespy (Garcia *et al.* 2018). A reference species tree was built using RAxML-ng (LG substitution model, 200 bootstraps) based on a MAFFT-linsi alignment of IF-2a proteins (identified as hits against the TIGR00491 HMM model). This tree recapitulates previously inferred relationships amongst the Methanobacteriales (Tokura *et al.* 2000).

Diversity at a given residue (column in the alignment) and for a given group (e.g. archaea with multiple histone paralogs) was calculated using the Shannon diversity index (*H*). Subsequently, we computed diversity ratios for two groups (A and B) as

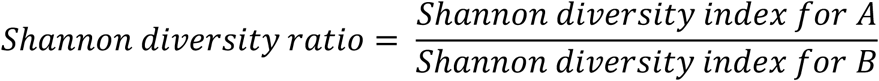

Similarity between histone groups in terms of the types of amino acids found at a given residue was calculated using the Jaccard index formula.

### Histone expression levels for different species

For *M. smithii, T. kodakarensis* and *T. onnurineus*, histone mRNA levels in exponential phase were obtained from NCBI’s Gene Expression Omnibus (GEO) and primary publications. The relative expression of histones in *T. kodakarensis* (Jäger *et al.* 2014) and *T. onnurineus* (GSE85760 (Cho *et al.* 2017)) is plotted as base mean and normalised mRNA respectively, in Figure S4. For *M. smithii*, we used the median value of histone expression across replicates for strain MsmPS in low formate, high hydrogen (GSE25408, (Hansen *et al.* 2011)).

### M. stadtmanae *culture, qRT-PCR analysis and proteomics*

*M. stadtmanae* DSM3091 was grown as previously described (Bang *et al.* 2012). Briefly, cultures were grown at 37°C in 50 ml minimal medium under strict anaerobic conditions. Medium was reduced with Na_2_S and cysteine (2 mM) and supplemented with 100 μg/ml ampicillin to prevent bacterial contamination. A 150 mM concentration of methanol and 1.5 atm H_2_-CO_2_ (80/20 [vol/vol]) served as carbon and energy source. Growth was monitored via turbidity at 600 nm (T_600_) and stopped at exponential or stationary phase by short incubation on ice (15 min) and subsequent centrifugation of cultures (3200× g for 30 min at 4°C). Resulting cell pellets were either resuspended in 500 *µ*l 50 mM TRIS containing RiboLock (Thermo Fisher Scientific) for RNA isolation or in 500 *µ*l 50 mM triethylammonium bicarbonate buffer for proteomics until further processing.

For both isolation of RNA and proteins, *M. stadtmanae* cells were lysed in liquid nitrogen using a Mikro-Dismembrator S laboratory ball mill (Sartorius) for 3 min at 1,600 bpm. For proteome analysis, cells were centrifuged after homogenisation at 15 700 × g and 4°C for 30 min and supernatant was used as cell-free protein extracts. RNA extraction and qRT-PCR assays were then performed as described earlier (Buddeweg *et al.* 2018). mRNA expression levels of three biological replicates were calculated using the normalizing 2-ΔΔCt value. *Msp_16S* and *Msp_rpoB* were used as genes for normalization (Dridi *et al.* 2009). Primers used are provided in Table S4.

Cell free protein extracts were run on a gel, low-molecular weight section (<10kDa) excised, and processed using a procedure adapted from (Shevchenko *et al.* 2006). Briefly, excised gel sections were further cut into cubes of approximately 2mm x 2mm and washed with 50mM ammonium bicarbonate in 50% aqueous acetonitrile (ACN). Dehydration of gel sections was carried out with 100% ACN. Sections were then sequentially reduced and alkylated with 10mM DTT and 55mM iodoacetamide, respectively. Digestions were carried out by addition of 500ng of trypsin per gel section, followed by incubation at 37°C overnight. Gel digest supernatants were then dried completely by vacuum centrifugation. Following extraction of tryptic peptides from gel pieces, dried extracts were reconstituted in 1% aqueous ACN, 0.1% formic acid (FA). Desalting was performed using C18 reverse phase solid phase extraction spin-tips (Glygen Corp.) following the manufacturer’s recommendations and eluted tryptic peptides were then dried by vacuum centrifugation.

Desalted gel digests were solubilised in 20*µ*l of 0.1% aqueous trifluoroacetic acid (TFA) and clarified solutions transferred to auto-sampler vials for LC-MS analysis. Peptides were separated using an Ultimate 3000 RSLC nano liquid chromatography system (Thermo Scientific) coupled to a LTQ Velos Orbitrap mass spectrometer (Thermo Scientific) via an EASY-Spray source. Sample aliquots (5.0 uL per injection) were loaded in technical duplicate onto a trapping column (Acclaim PepMap 100 C18, 100μm × 2cm) at 8μL/min in 2% ACN, 0.1% TFA. Peptides were then eluted on-line to an analytical column (EASY-Spray PepMap C18, 75μm × 25cm) and peptides were separated using a stepped 90 minute gradient: 4-25% of buffer B for 60 minutes, 25-45% buffer B for 30 minutes. Buffer compositions were buffer A: 2% ACN, 0.1% FA; buffer B: 80% ACN, 0.1% FA. Eluted peptides were analysed by the LTQ Velos operating in positive ion polarity using a data-dependent acquisition mode. Ions were selected for fragmentation from an initial MS1 survey scan at 15,000 resolution (at m/z 200), followed by Ion Trap collisional induced dissociation (CID) of the top 10 most abundant ions. MS1 and MS2 scan automatic gain control (AGC) targets were set to 1e6 and 1e4 for a maximum injection times of 500ms and 100ms respectively. A survey scan with m/z range of 350 – 1500 was used, with a normalised collision energy (NCE) set to 35%, charge state rejection enabled for +1 ions and a minimum threshold for triggering fragmentation of 500 counts.

The resulting data were processed using the MaxQuant software platform (v1.5.3.8), with database searches carried out by the in-built Andromeda search engine against the *M. stadtmanae* DSM3091 proteome as annotated in NCBI. A reverse decoy database search approach was used at a 1% false discovery rate (FDR) for peptide spectrum matches and protein identifications. Search parameters included: maximum missed cleavages set to 2, fixed modification of cysteine carbamidomethylation and variable modifications of methionine oxidation, protein N-terminal and lysine acetylation, glutamine to pyro-glutamate conversion, asparagine deamidation as well as lysine and arginine methylation. Label-free quantification was enabled with an LFQ minimum ratio count of 2. The ‘match between runs’ function was used with match and alignment time settings of 0.7 and 20 minutes respectively.

## ACKNOWLEDGEMENTS

We thank Samuel Bowerman, Remus Dame, and Karolin Luger for feedback on the manuscript, members of the Molecular Systems and DNA Replication groups for discussion, and Pierre Garcia for advice on phylogenetic tools. This work was funded by Medical Research Council core funding (TW), a Medical Research Council studentship (KMS), a UKRI Innovation Fellowship (JBS), EMBO Short-Term Fellowship 8472 (AH) and DFG grant SCHM1052/11-2 (RAS). SG acknowledges funding from the French National Agency for Research, Grant ArchEvol (ANR-16-CE02-0005-01). This project made use of time on UK Tier 2 JADE granted via the UK High-End Computing Consortium for Biomolecular Simulation, HECBioSim, supported by the EPSRC (grant no. EP/R029407/1).

## SUPPLEMENTARY FIGURE LEGENDS

**Figure S1.**
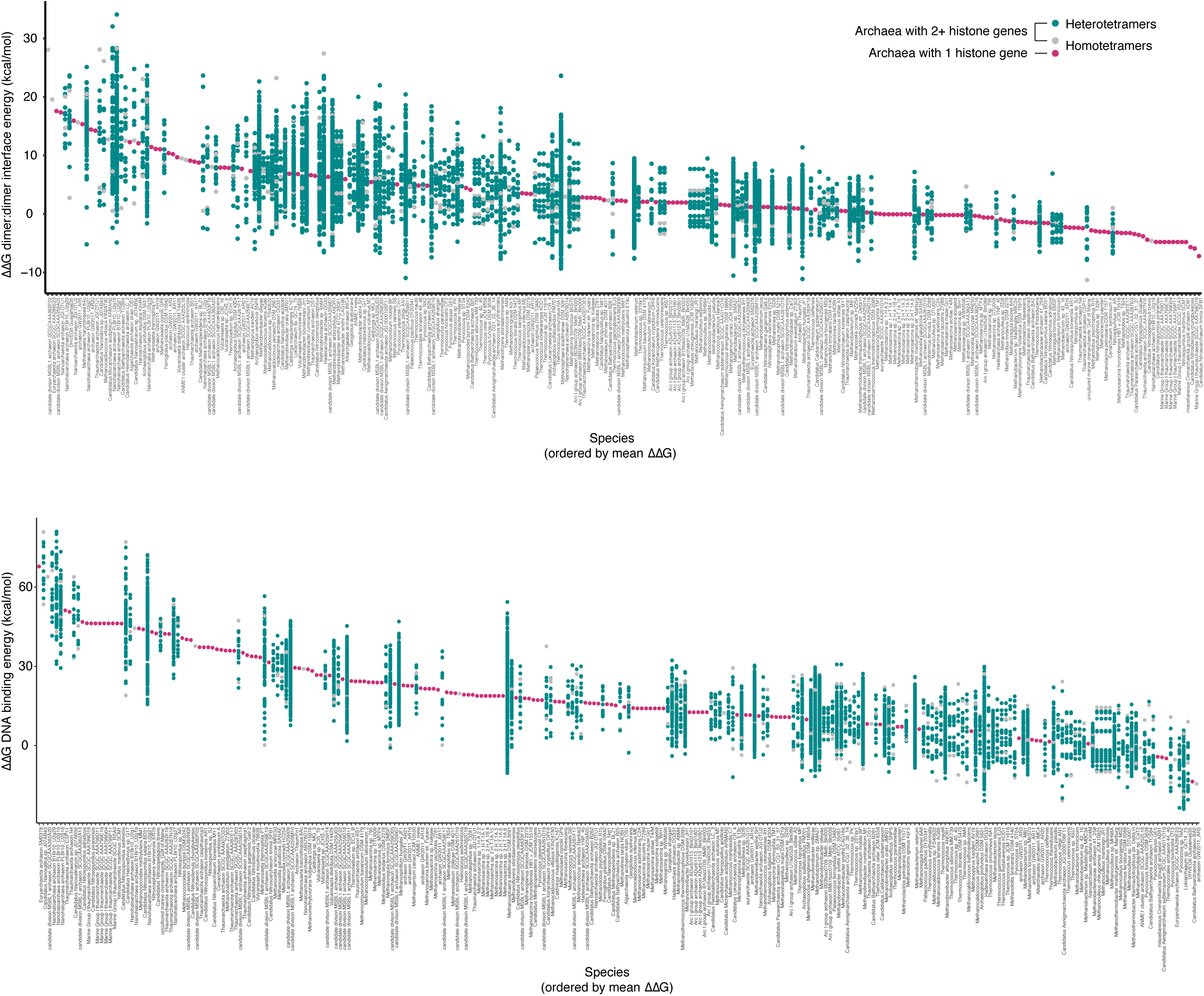
DNA binding strength and tetramerization strength (dimer:dimer interface energy) for every possible tetrameric histone complex within each species of archaea in our sample. Each point, grouped by species, represents an individual complex. Species are ordered by mean interaction energy across tetramers. ΔΔG is relative to HMfA.

**Figure S2.**
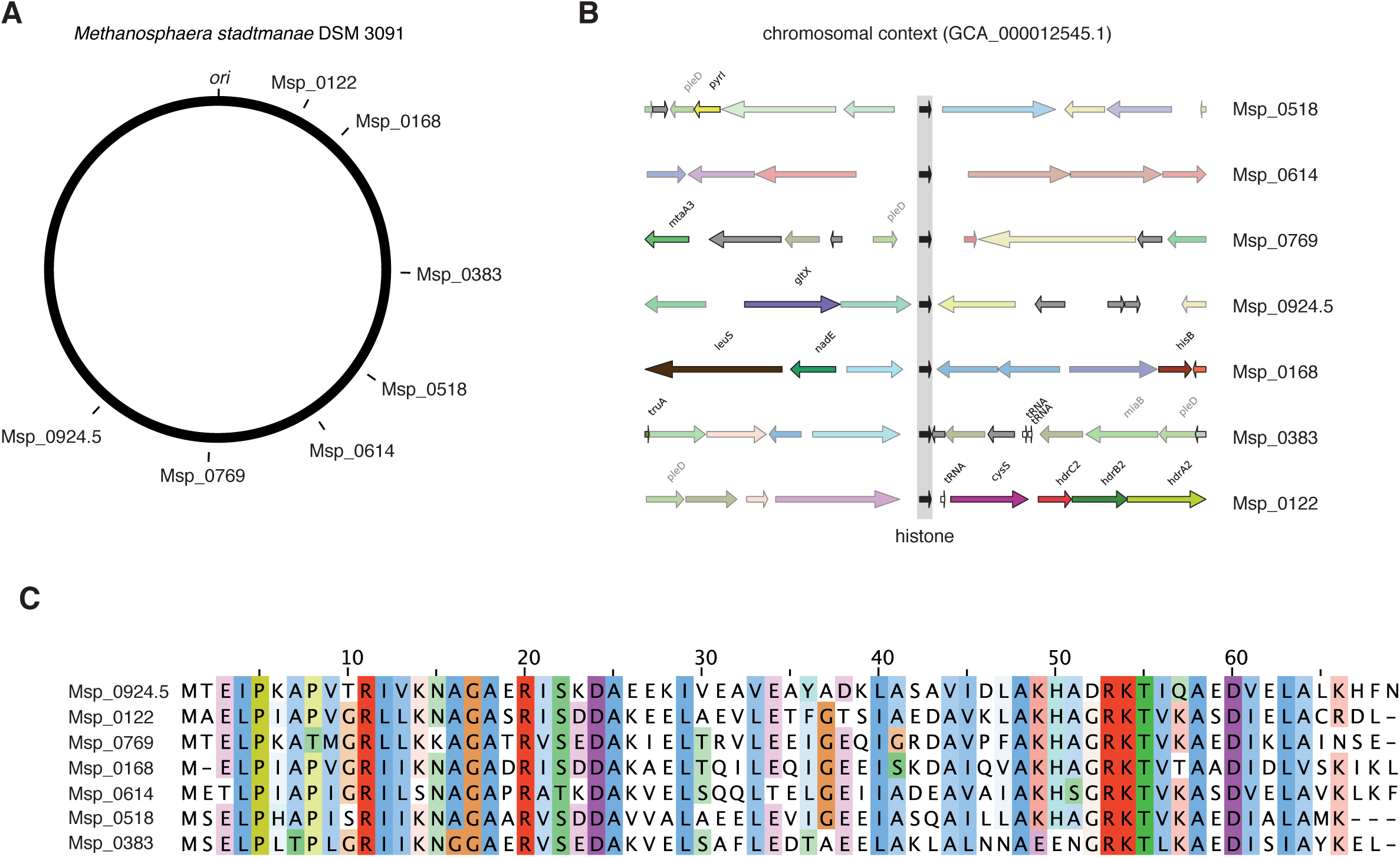
(**A**) Circular chromosome map of *M. stadtmanae* DSM 3091 showing the locations of the seven histone paralogs. Note that one of the histone paralogs is not annotated in GenBank. As it is located between Msp_0924 and Msp_0925, we have labelled it Msp_0924.5. (**B**) Chromosomal context of *M. stadtmanae* histone paralogs. Histones are flanked by comparatively large intergenic regions upstream and either large intergenic regions or convergently transcribed genes downstream, suggesting that histones are transcribed as single-gene operons. (**C**) Protein-level alignment of the seven paralogs.

**Figure S3.**
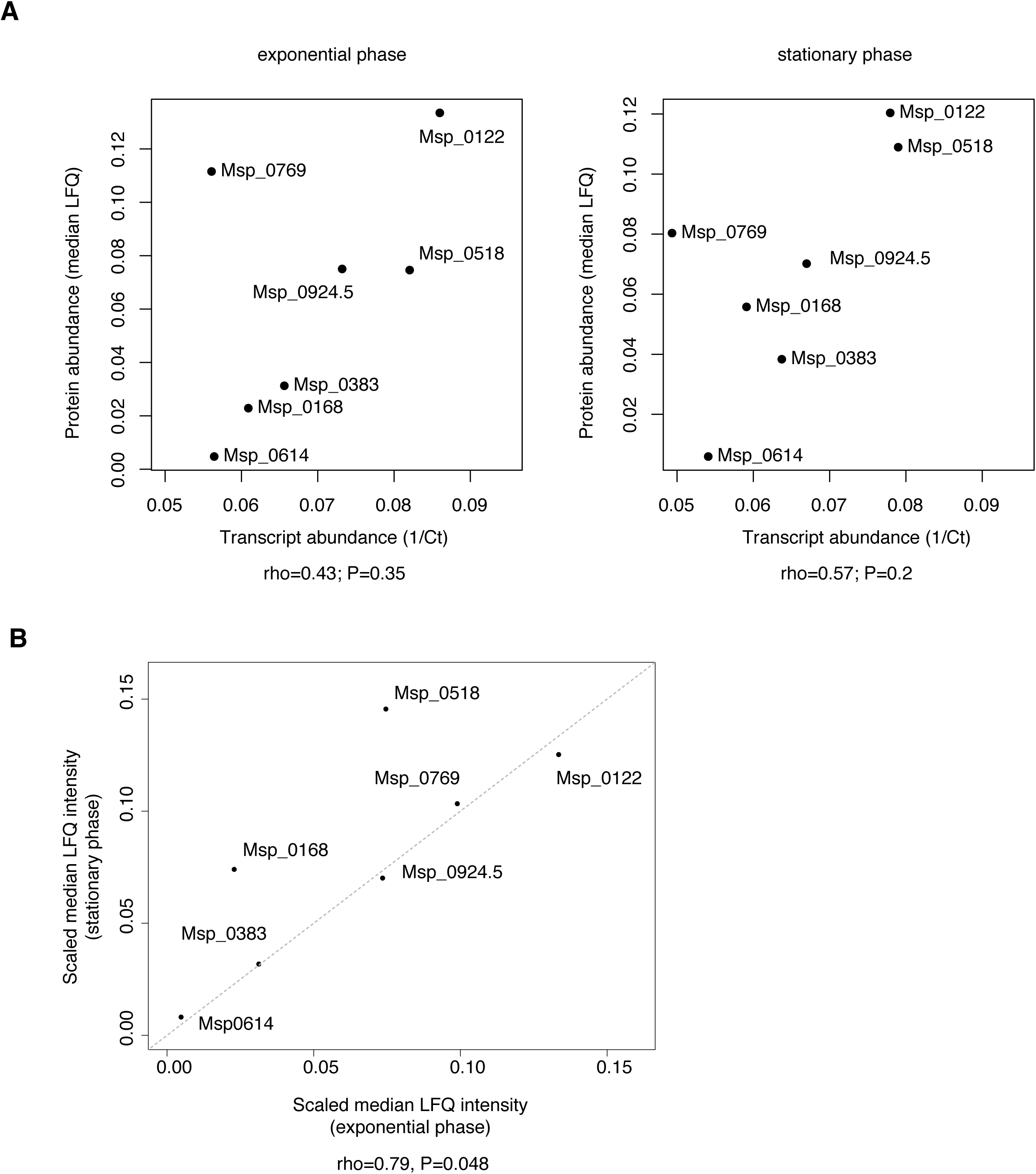
(**A**) Protein abundance and transcript abundance for *M. stadtmanae* histone paralogs in exponential (left) and stationary (right) phase. (**B**) Correlation between median protein abundance (LFQ intensity) for *M. stadtmanae* histone paralogs in exponential versus stationary phase.

**Figure S4.**
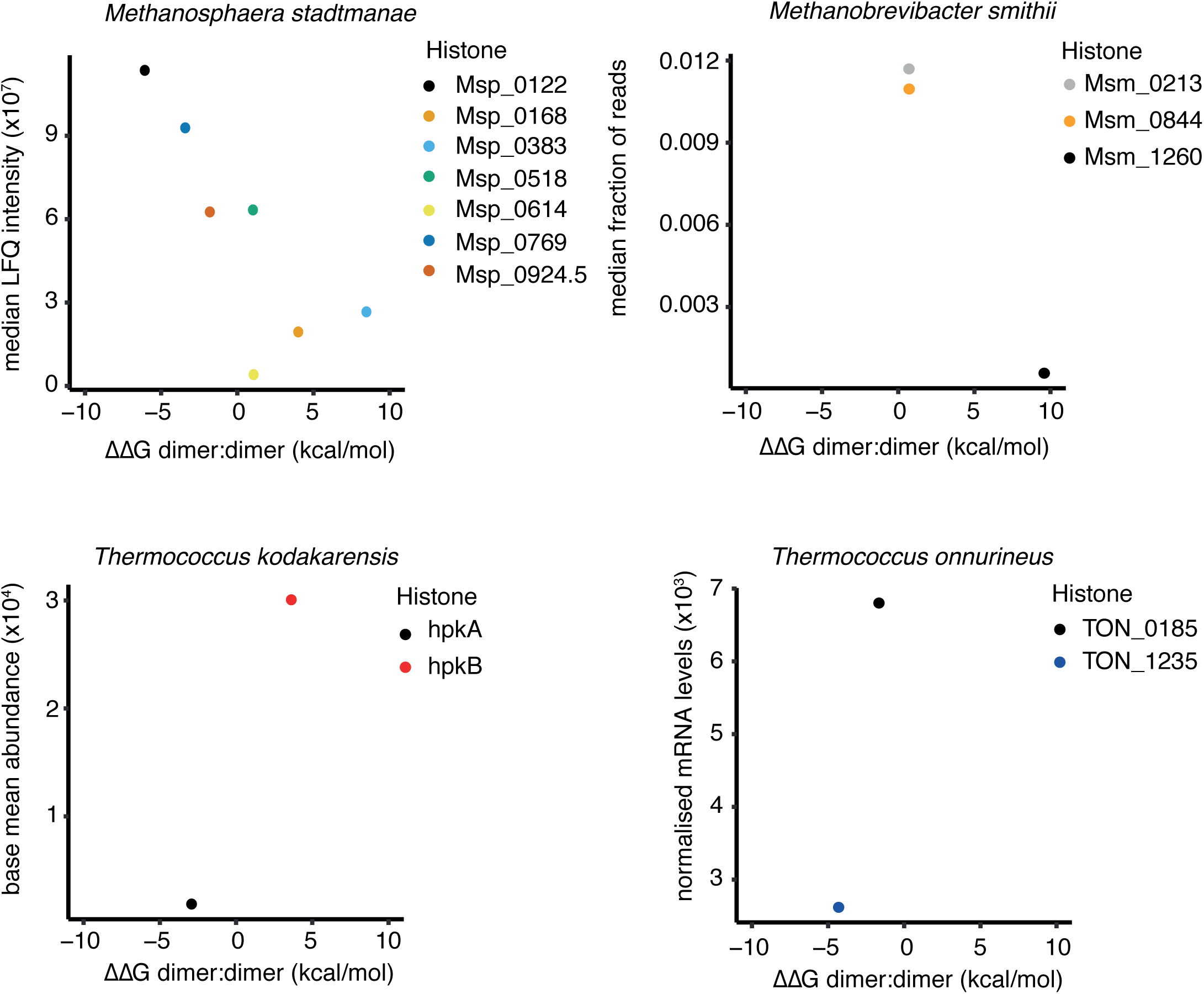
Tetramerisation strength for homotetrameric histone models in *M. stadtmanae, M. smithii, T. kodakarensis* and *T. onnurineus* and its relation to paralog expression levels as measured by protein or transcript abundance in exponential phase. ΔΔG is relative to HMfA. See Methods for data provenance.

**Figure S5.**
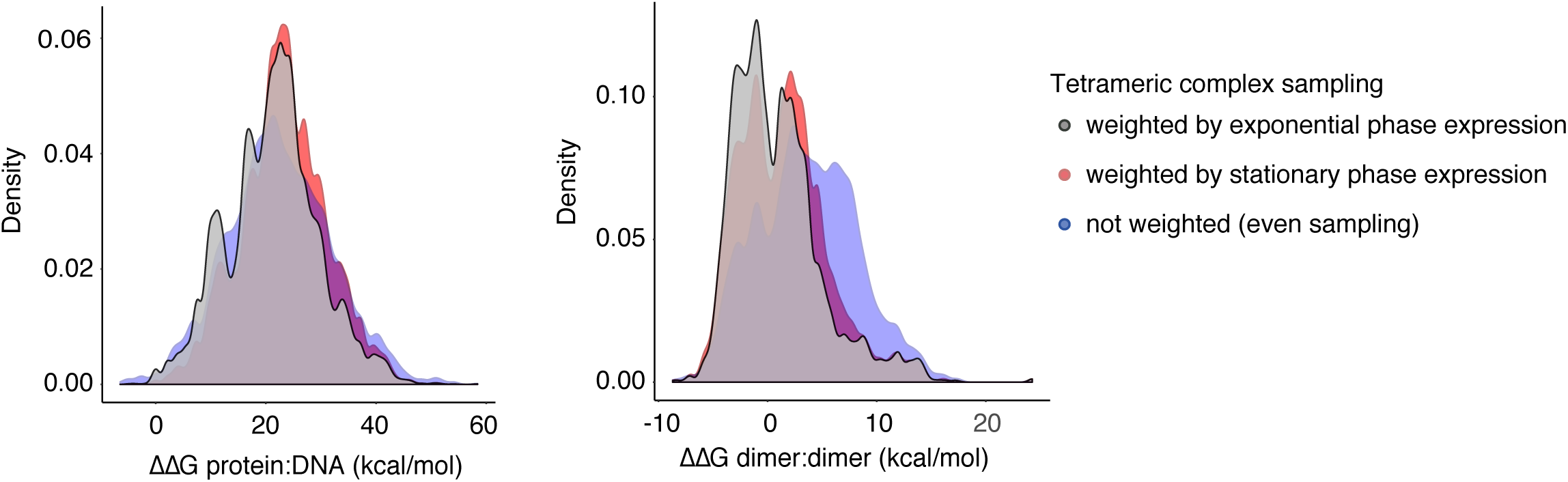
Distribution of DNA binding and tetramerization strength for 100,000 histone tetrameric complexes in *M. stadtmanae*, with tetramer composition determined by sampling based on relative paralog abundance (mean LFQ intensity) in exponential phase, stationary phase, and random sampling.

**Figure S6.**
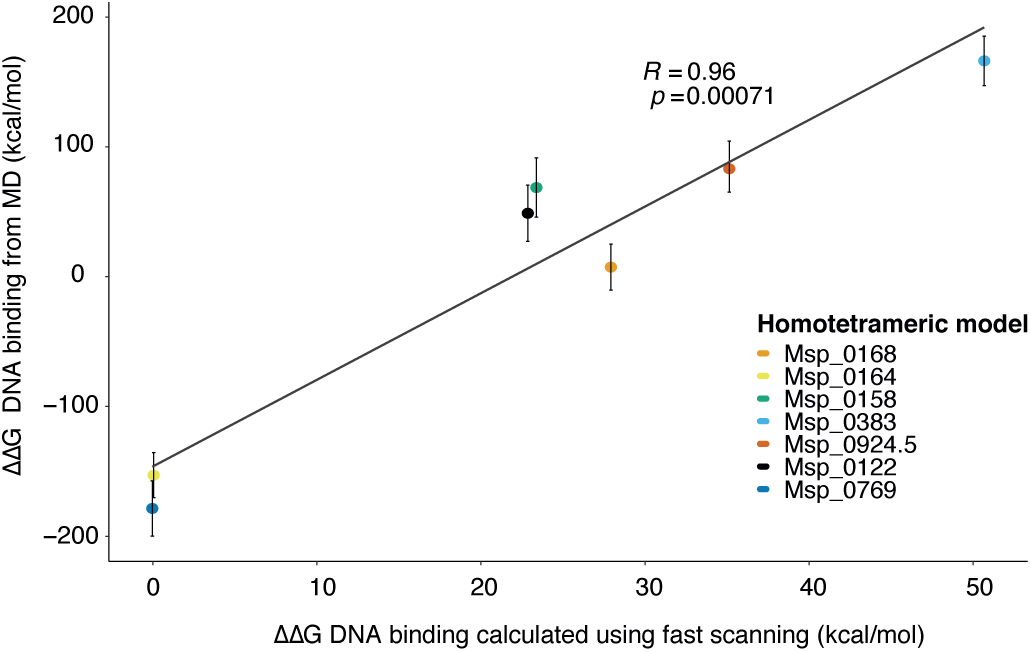
DNA binding (ΔΔG relative to Msp_0769) across 100 ns of MD simulations compared to fast mutational scanning results for homotetrameric histone complexes from *M. stadtmanae*.

**Figure S7.**
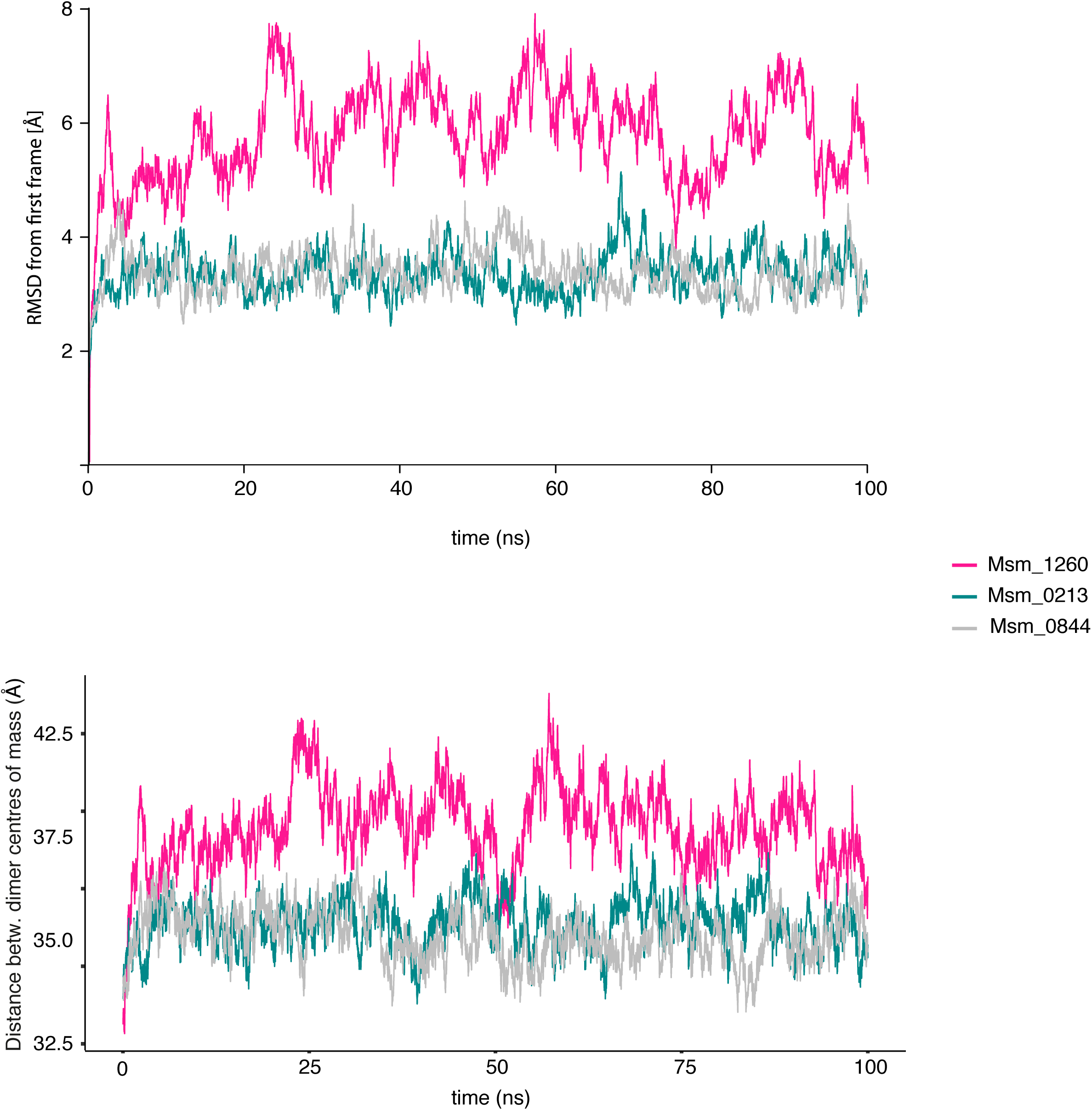
MD simulations of homotetrameric histone-DNA complexes for paralogs from *M. smithii* showing (top) RMSD and (bottom) distance between centre of mass of each dimer over 100 ns.

**Figure S8.**
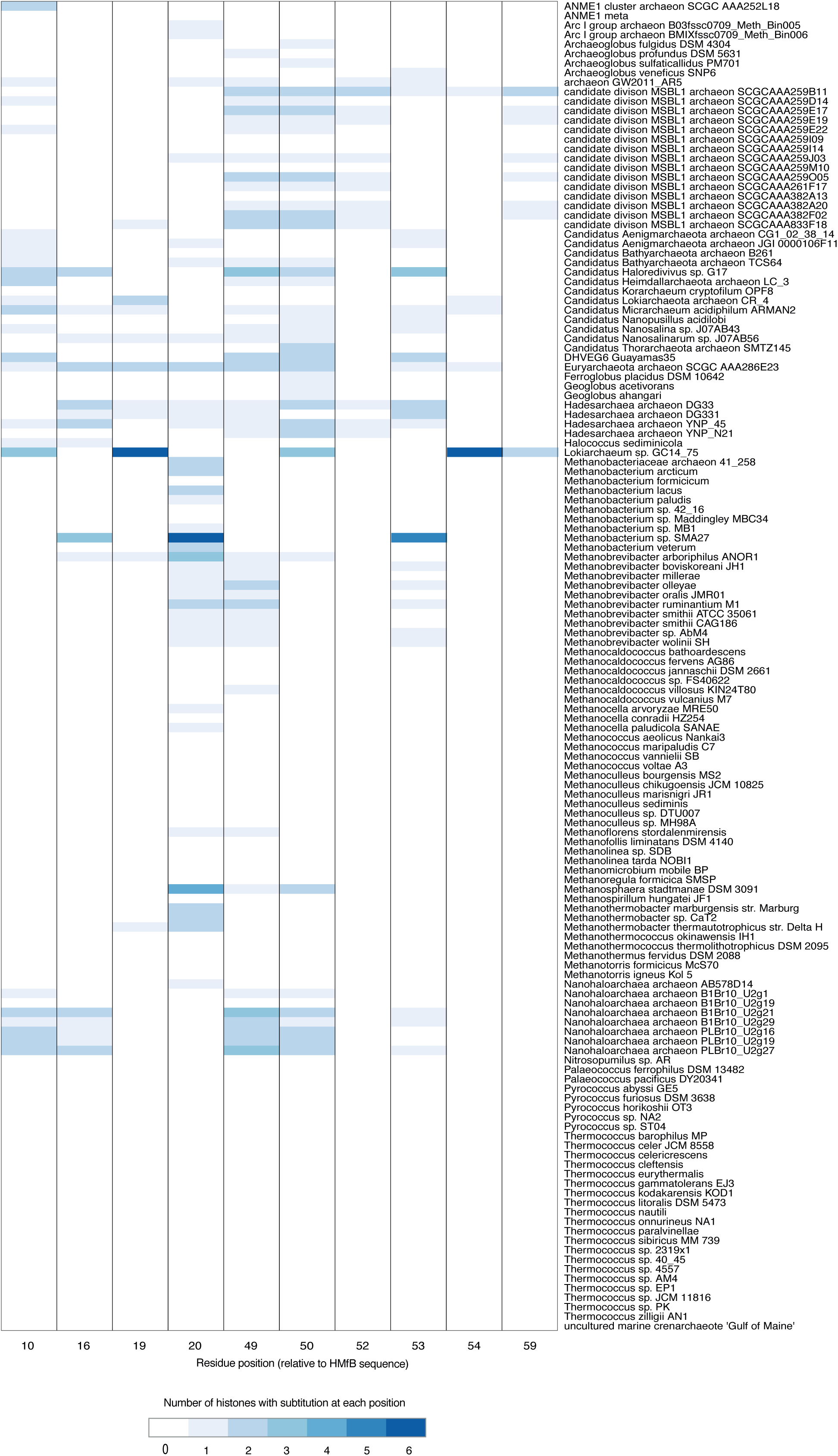
Distribution of amino acid diversity at selected structurally sensitive sites in archaea with more than one histone. Substitutions are defined as amino acids which differ from the most commonly found amino acid at that position. Species are ordered alphabetically.

**Figure S9.**
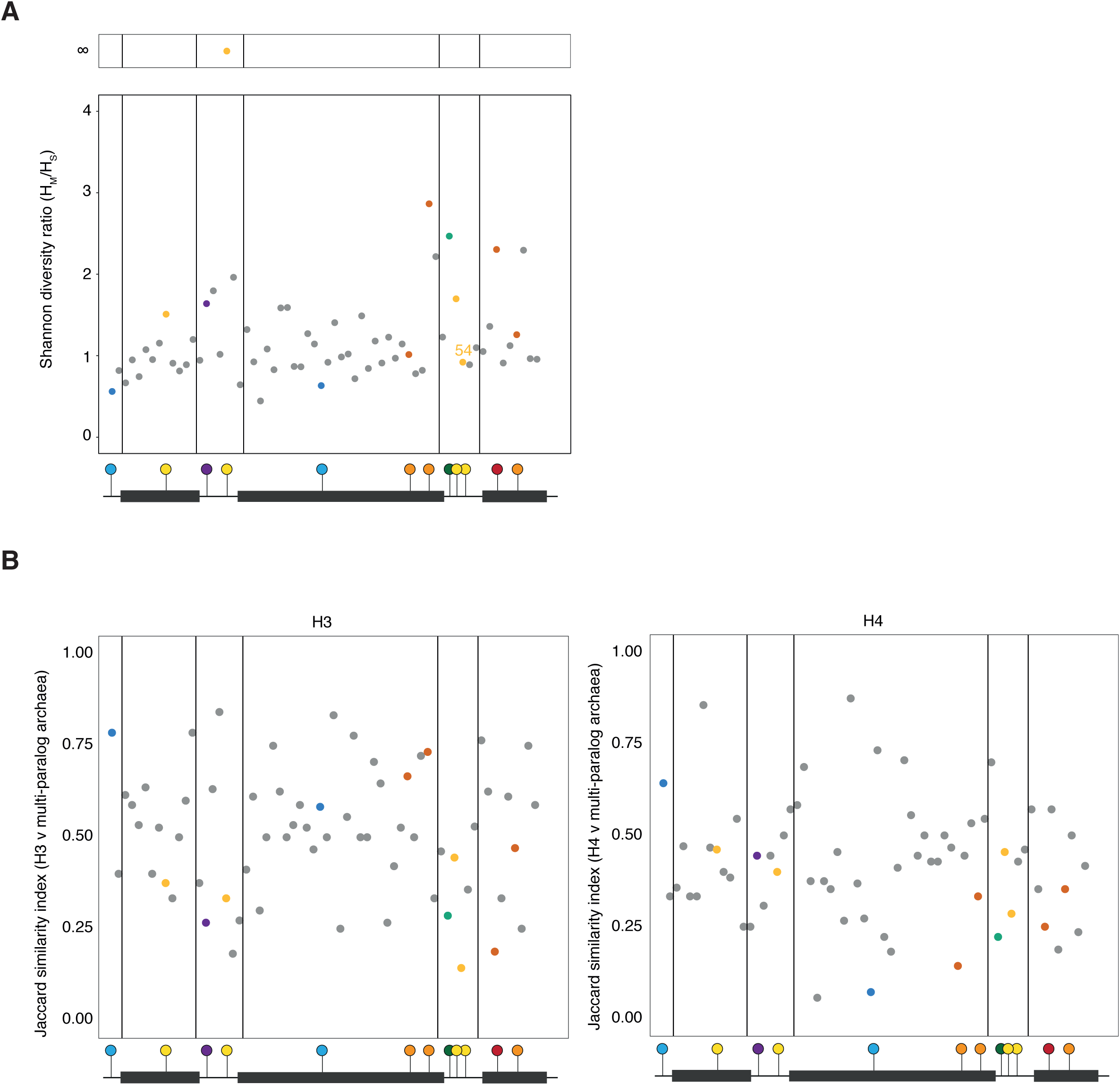
(**A**) Shannon diversity ratio (*H*_*M*_*/H*_*S*_) across the histone fold excluding Asgard archaea. Note the precipitous drop in the Shannon ratio at residue 54 compared to Figure 7. (**B**) Jaccard similarity at each position in the core histone fold domain comparing archaea with more than one histone gene to H3 (left) and H4 (right).

**Figure S10.**
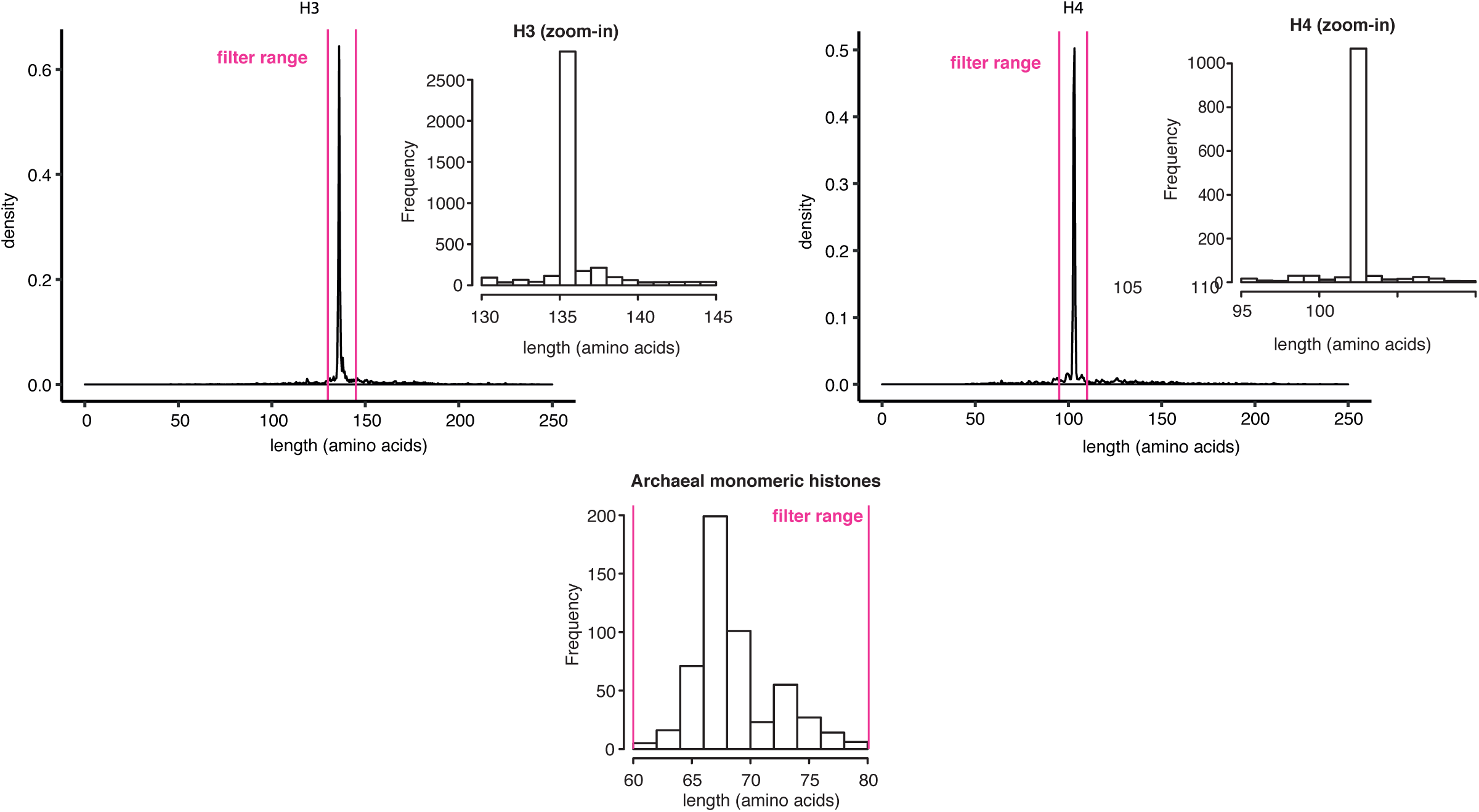
Length distribution of putative H3 and H4 proteins prior to filtering. Size thresholds applied are highlighted in pink. The distribution of histone lengths in archaea after filtering is given below.

## SUPPLEMENTARY DATA

**Table S1.** Tetramerization and DNA binding energies for all homo-an heterotetrameric histone-DNA complexes in 282 archaea.

**Table S2.** Predicted impact of single amino acid substitutions in HMfB compared to the study of Soares *et al.* (2000)

**Table S3.** Residue numbering for *M. fervidus* HMfB and *S. cerevisiae* H3 (P61830) and H4 (P02309).

**Table S4.** Primers used for qRT-PCR.

**File S1.** Video of molecular dynamics simulation trace for the Msp_0383 tetramer.

**File S2.** Video of molecular dynamics simulation trace for the Msp_0769 tetramer.

**File S3.** Alignment of IF-2A orthologs.

**File S4.** Alignment of 560 archaeal histones.

**File S5.** Alignment of augmented set of 168 Methanobacteriales histones (including additional *M. stadtmanae* genomes).

**File S6.** Reference phylogeny based on alignment in File S3. Bootstrap values are given as a percentage of 200 non-parametric bootstraps.

**File S7.** Pan-archaeal phylogeny of histones, reconstructed based on alignment in File S4. Bootstrap values are given as a percentage of 1000 ultra-fast bootstraps.

**File S8.** Phylogeny of histones in the Methanobacteriales, reconstructed based on alignment in File S5. Bootstrap values are given as a percentage of 500 non-parametric bootstraps.

---

**Table S2.** HMfB single amino acid mutants characterised in gel shift assays (Soares et al. 2000)

| Mutation | $\Delta\Delta G$ DNA binding<br>(kcal/mol) | $\Delta\Delta G$ stability of the<br>protein (kcal/mol) | Gel shift result from<br>Soares *et al.* (2000) |
| --- | --- | --- | --- |
| M1G | -0.229 | 2.373 | Same gel shift as WT |
| E2D | -1.5081 | 0.725 | Decreased gel shift |
| E2K | -1.8864 | -17.952 | Same gel shift |
| L3C | -0.866 | 12.1636 | Did not accumulate in E. coli |
| L3I | -0.4955 | 0.234 | Decreased gel shift |
| P4C | -0.7506 | 15.609 | Same gel shift |
| P4S | 3.5532 | 14.2371 | Same gel shift |
| I5V | -1.0162 | 3.758 | Same gel shift |
| A6P | 30.558863 | -3.104 | Did not accumulate in E. coli |
| P7A | 1.4649 | 13.8786 | Same gel shift |
| I8V | 0.0469 | 4.354 | Same gel shift |
| G9D | 0.9748 | 0.818 | Did not accumulate in E. coli |
| R10S | 1.41 | 0.315 | Did not accumulate in E. coli |
| R10K | 1.8425 | -6.518 | Decreased gel shift |
| R10G | 1.4124 | 0.416 | Did not accumulate in E. coli |
| I11L | -0.7244 | -4.562 | Same gel shift |
| K13E | -0.2283 | -4.133 | Did not accumulate in E. coli |
| K13Q | -0.4862 | -1.443 | Did not accumulate in E. coli |
| K13T | -0.9661 | 4.569 | Did not accumulate in E. coli |
| K13R | -2.0568 | -1.117 | Same gel shift |
| D14E | -0.3723 | -5.831 | Did not accumulate in E. coli |
| D14N | -0.493 | -9.57 | Same gel shift |
| D14K | -1.24 | -16.484 | Decreased gel shift |
| D14H | -3.0588 | -14.311 | Did not accumulate in E. coli |
| A15S | 0 | 7.3696 | Same gel shift |
| A15G | 0 | 7.3371 | Did not accumulate in E. coli |
| E18D | 1.2779 | 0.034 | Same gel shift |
| E18P | 0.4728 | -5.265 | Same gel shift |
| E18K | -1.4242 | 3.089 | Did not accumulate in E. coli |
| R19I | 4.3091 | -3.551 | Did not accumulate in E. coli |
| R19Q | 5.2768 | -2.048 | Did not accumulate in E. coli |
| R19S | 9.9084 | 1.502 | Did not accumulate in E. coli |
| R19G | 10.5705 | 5.8858 | Decreased gel shift |
| R19K | 5.6178 | -1.358 | Decreased gel shift |
| V20C | -0.4124 | 9.6386 | Did not accumulate in E. coli |
| V20I | 0.3076 | -6.426 | Same gel shift |
| V20D | -1.2659 | 19.0681 | Did not accumulate in E. coli |
| S21A | 1.9393 | 1.831 | Decreased gel shift |
| S21C | -1.3203 | 7.5993 | Same gel shift |
| S21T | 1.5381 | -1.531 | Decreased gel shift |
| R25K | 5.5543 | -6.126 | Same gel shift |
| I26E | 0.3859 | 5.25 | Decreased gel shift |
| T27A | -0.0002 | 3.174 | Same gel shift |
| T27C | 0 | 6.5756 | Same gel shift |
| I31V | 0.0005 | 6.4488 | Same gel shift |
| I31C | -0.0019 | 15.9234 | Same gel shift |
| L32C | 0 | 19.2103 | Did not accumulate in E. coli |
| M35C | 0 | 10.6139 | Same gel shift |
| R37K | -0.1871 | -8.6 | Same gel shift |
| R37L | 0.024 | -4.509 | Decreased gel shift |
| R37Q | -0.014 | 2.771 | Decreased gel shift |
| D38E | 0 | -3.054 | Same gel shift |
| I39V | 0 | 1.634 | Same gel shift |
| I39C | 0 | 11.2149 | Same gel shift |
| A43I | 0.0061 | 32.1119 | Did not accumulate in E. coli |
| S41C | 0.012 | 0.715 | Same gel shift |
| I44V | 0 | 3.902 | Same gel shift |
| L46I | 0.0158 | 6.3608 | Decreased gel shift |
| L46V | 0.0079 | 10.9782 | Decreased gel shift |
| L46F | -0.0037 | 4.347 | Same gel shift |
| L46S | 0.0021 | 20.4698 | Did not accumulate in E. coli |
| R48K | 0.3128 | -0.278 | Same gel shift |
| H49A | 0.1917 | 1.99 | Decreased gel shift |
| H49D | 0.5219 | 11.2511 | Same gel shift |
| G51K | -0.9888 | 8.6454 | Did not accumulate in E. coli |
| G51A | -0.677 | 11.4035 | Did not accumulate in E. coli |
| R52K | -0.252 | 1.341 | Decreased gel shift |
| R52A | 1.4069 | 5.407 | Did not accumulate in E. coli |
| R52H | 4.0269 | 5.6672 | Did not accumulate in E. coli |
| R52Q | -0.5464 | 3.51 | Did not accumulate in E. coli |
| K53R | 1.6919 | -1.096 | Decreased gel shift |
| K53E | 3.2454 | 1.266 | Did not accumulate in E. coli |
| K53T | 2.1058 | 2.286 | Did not accumulate in E. coli |
| T54A | 3.5459 | 2.269 | Did not accumulate in E. coli |
| T54C | 2.7237 | 1.158 | Did not accumulate in E. coli |
| T54K | 4.9682 | 1.889 | Did not accumulate in E. coli |
| T54R | 7.0644 | 4.545 | Did not accumulate in E. coli |
| T54S | 2.2691 | 4.55 | Did not accumulate in E. coli |
| T54V | -1.0832 | -2.153 | Did not accumulate in E. coli |
| T54Y | 19.2299 | 19.3918 | Did not accumulate in E. coli |
| I55V | 0.662 | 4.123 | Same gel shift |
| I55L | 0.2032 | -0.262 | Decreased gel shift |
| I55T | 0.6911 | 17.92 | Decreased gel shift |
| I55M | -0.2553 | 4.661 | Decreased gel shift |
| I55C | 0.285 | 16.4504 | Did not accumulate in E. coli |
| K56R | -1.1339 | 0.905 | Same gel shift |
| K56T | -1.529 | 2.93 | Did not accumulate in E. coli |
| K56E | 2.0138 | -1.73 | Did not accumulate in E. coli |
| K56I | -0.6459 | 4.121 | Did not accumulate in E. coli |
| K56Q | -1.3688 | 0.836 | Did not accumulate in E. coli |
| E58S | -1.7313 | 5.415 | Did not accumulate in E. coli |
| D59A | 1.3352 | 9.3873 | Did not accumulate in E. coli |
| D59E | -1.5257 | 5.7071 | Did not accumulate in E. coli |
| D59N | -0.1062 | 13.877 | Did not accumulate in E. coli |
| I60V | 0 | 4.055 | Same gel shift |
| E61A | 0.006 | 1.404 | Same gel shift |
| E61V | -0.0047 | 3.984 | Same gel shift |
| E61Q | 0.0064 | -1.091 | Same gel shift |
| E61K | 0.0129 | -2.887 | Decreased gel shift |
| E61R | -0.0027 | -2.443 | Decreased gel shift |
| L62I | 0.0099 | 5.129 | Decreased gel shift |
| L62V | 0.007 | 7.5351 | Decreased gel shift |
| L62Y | -0.096 | 3.525 | Same gel shift |
| L62M | 0.0173 | -2.076 | Did not accumulate in E. coli |
| V64R | -0.0057 | 4.251 | Decreased gel shift |
| R65K | -0.0062 | 2.925 | Same gel shift |
| R66M | 0.0066 | -1.184 | Same gel shift |

**Table S3.** HMfB and *S. cerevisiae* H3/H4 sequences and residue numbering for each position in the alignment used for diversity analysis

| *M. fervidus* HMfB residue number | *M. fervidus* HMfB residue | *S. cerevisiae* H3 residue number (P61830) | *S. cerevisiae* H3 residue | *S. cerevisiae* H4 residue number (P02309) | *S. cerevisiae* H4 residue |
| --- | --- | --- | --- | --- | --- |
| 2 | E | 60 | E | 29 | G |
| 3 | L | 63 | I | 30 | I |
| 4 | P | 64 | R | 31 | T |
| 5 | I | 65 | K | 32 | K |
| 6 | A | 66 | L | 33 | P |
| 7 | P | 67 | P | 34 | A |
| 8 | I | 68 | F | 35 | I |
| 9 | G | 69 | Q | 36 | R |
| 10 | R | 70 | R | 37 | R |
| 11 | I | 71 | L | 38 | L |
| 12 | I | 72 | V | 39 | A |
| 13 | K | 73 | R | 40 | R |
| 14 | D | 74 | E | 41 | R |
| 15 | A | 75 | I | 42 | G |
| 16 | G | 76 | A | 43 | G |
| 17 | A | 77 | Q | 44 | V |
| 18 | E | 78 | D | 45 | K |
| 19 | R | 84 | R | 46 | R |
| 20 | V | 85 | F | 47 | I |
| 21 | S | 86 | Q | 48 | S |
| 22 | D | 87 | S | 49 | G |
| 23 | D | 88 | S | 50 | L |
| 24 | A | 89 | A | 51 | I |
| 25 | R | 90 | I | 52 | Y |
| 26 | I | 91 | G | 53 | E |
| 27 | T | 92 | A | 54 | E |
| 28 | L | 93 | L | 55 | V |
| 29 | A | 94 | Q | 56 | R |
| 30 | K | 95 | E | 57 | A |
| 31 | I | 96 | S | 58 | V |
| 32 | L | 97 | V | 59 | L |
| 33 | E | 98 | E | 60 | K |
| 34 | E | 99 | A | 61 | S |
| 35 | M | 100 | Y | 62 | F |
| 36 | G | 101 | L | 63 | L |
| 37 | R | 102 | V | 64 | E |
| 38 | D | 103 | S | 65 | S |
| 39 | I | 104 | L | 66 | V |
| 40 | A | 105 | F | 67 | I |
| 41 | S | 106 | E | 68 | R |
| 42 | E | 107 | D | 69 | D |
| 43 | A | 108 | T | 70 | S |
| 44 | I | 109 | N | 71 | V |
| 45 | K | 110 | L | 72 | T |
| 46 | L | 111 | A | 73 | Y |
| 47 | A | 112 | A | 74 | T |
| 48 | R | 113 | I | 75 | E |
| 49 | H | 114 | H | 76 | H |
| 50 | A | 115 | A | 77 | A |
| 51 | G | 116 | K | 78 | K |
| 52 | R | 117 | R | 79 | R |
| 53 | K | 118 | V | 80 | K |
| 54 | T | 119 | T | 81 | T |
| 55 | I | 120 | I | 82 | V |
| 56 | K | 121 | Q | 83 | T |
| 57 | A | 122 | K | 84 | S |
| 58 | E | 123 | K | 85 | L |
| 59 | D | 124 | D | 86 | D |
| 60 | I | 125 | I | 87 | V |
| 61 | E | 126 | K | 88 | V |
| 62 | L | 127 | L | 89 | Y |
| 63 | A | 128 | A | 90 | A |
| 64 | V | 129 | R | 91 | L |
| 65 | R | 130 | R | 92 | K |

**Table S4.**
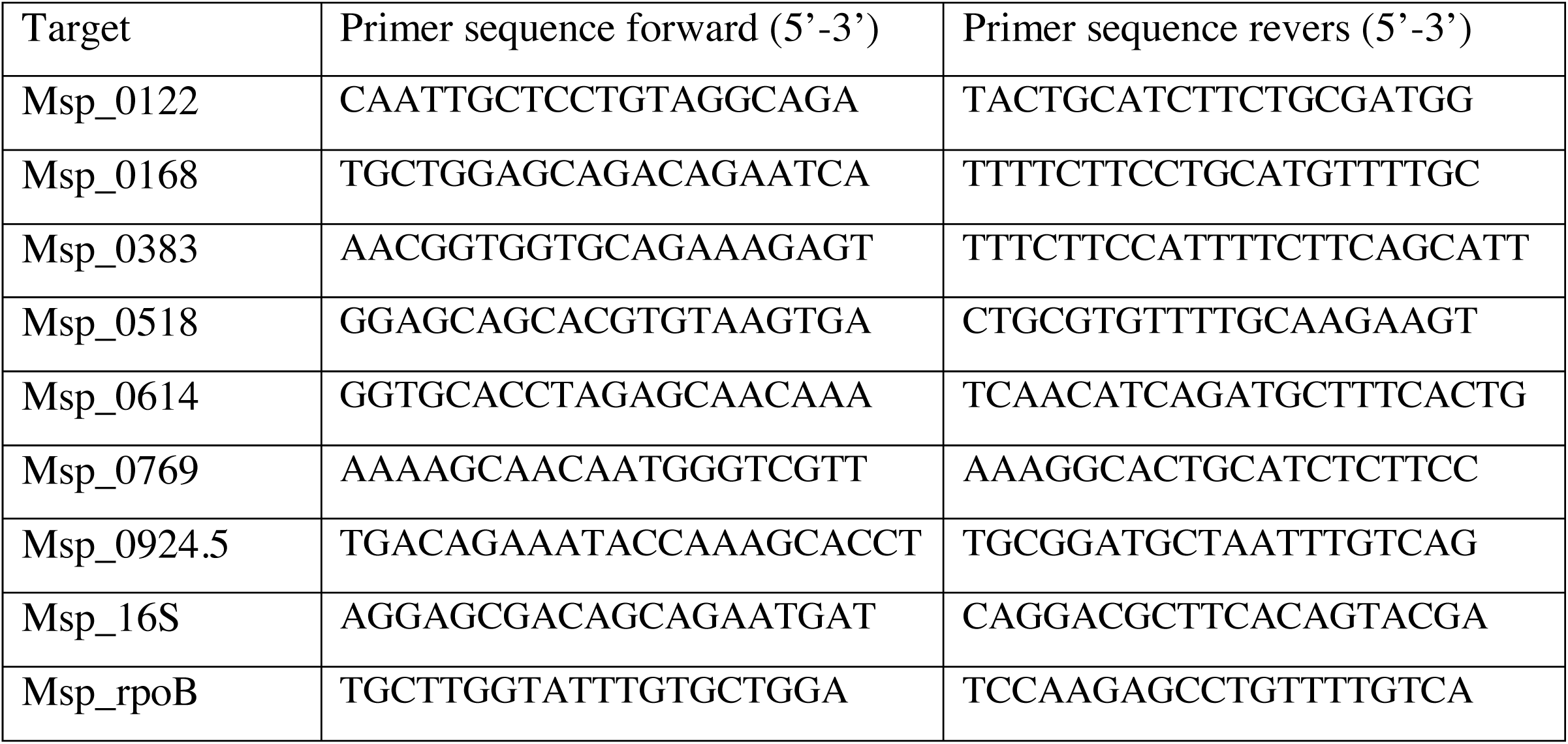
Primers used for qRT-PCR.

## REFERENCES

Adam P. S., Borrel G., Brochier-Armanet C., Gribaldo S., 2017 The growing tree of Archaea: new perspectives on their diversity, evolution and ecology. ISME J 11: 2407–2425.

Bang C., Schilhabel A., Weidenbach K., Kopp A., Goldmann T., Gutsmann T., Schmitz R. A., 2012 Effects of antimicrobial peptides on methanogenic archaea. Antimicrob. Agents Chemother. 56: 4123–4130.

Berendsen H. J. C., Postma J. P. M., van Gunsteren W. F., DiNola A., Haak J. R., 1998 Molecular dynamics with coupling to an external bath. The Journal of Chemical Physics 81: 3684–3690.

Betts H. C., Puttick M. N., Clark J. W., Williams T. A., Donoghue P. C. J., Pisani D., 2018 Integrated genomic and fossil evidence illuminates life’s early evolution and eukaryote origin. Nat Ecol Evol 2: 1556–1562.

Brunk C. F., Martin W. F., 2019 Archaeal Histone Contributions to the Origin of Eukaryotes. Trends in Microbiology 27: 703–714.

Buddeweg A., Sharma K., Urlaub H., Schmitz R. A., 2018 sRNA41 affects ribosome binding sites within polycistronic mRNAs in Methanosarcina mazei Gö1. Molecular Microbiology 107: 595–609.

Cho S., Kim M.-S., Jeong Y., Lee B.-R., Lee J.-H., Kang S. G., Cho B.-K., 2017 Genomewide primary transcriptome analysis of H 2 -producing archaeon Thermococcus onnurineus NA1. Scientific Reports 7: 1–12.

Cubonova L., Katano M., Kanai T., Atomi H., Reeve J. N., Santangelo T. J., 2012 An Archaeal Histone Is Required for Transformation of Thermococcus kodakarensis. Journal of Bacteriology 194: 6864–6874.

Davidchack R. L., Handel R., Tretyakov M. V., 2009 Langevin thermostat for rigid body dynamics. The Journal of Chemical Physics 130: 234101.

Decanniere K., Babu A. M., Sandman K., Reeve J. N., Heinemann U., 2000 Crystal structures of recombinant histones HMfA and HMfB from the hyperthermophilic archaeon Methanothermus fervidus. Journal of Molecular Biology 303: 35–47.

Dridi B., Henry M., Khéchine El A., Raoult D., Drancourt M., 2009 High Prevalence of Methanobrevibacter smithii and Methanosphaera stadtmanae Detected in the Human Gut Using an Improved DNA Detection Protocol. PLoS ONE 4: e7063.

Eme L., Sharpe S. C., Brown M. W., Roger A. J., 2014 On the age of eukaryotes: evaluating evidence from fossils and molecular clocks. Cold Spring Harbor Perspectives in Biology 6: a016139.

Eme L., Spang A., Lombard J., Stairs C. W., Ettema T. J. G., 2017 Archaea and the origin of eukaryotes. Nature Reviews Microbiology 15: 711–723.

Essmann U., Perera L., Berkowitz M. L., Darden T., Lee H., Pedersen L. G., 1995 A smooth particle mesh Ewald method. The Journal of Chemical Physics 103: 8577–8593.

Filipescu D., Müller S., Almouzni G., 2014 Histone H3 variants and their chaperones during development and disease: contributing to epigenetic control. Annu Rev Cell Dev Biol 30: 615–646.

Friedrich-Jahn U., Aigner J., Längst G., Reeve J. N., Huber H., 2009 Nanoarchaeal origin of histone H3? Journal of Bacteriology 191: 1092–1096.

Garcia P. S., Jauffrit F., Grangeasse C., Brochier-Armanet C., 2018 GeneSpy, a user-friendly and flexible genomic context visualizer. Bioinformatics 35: 329–331.

Götz A. W., Williamson M. J., Xu D., Poole D., Le Grand S., Walker R. C., 2012 Routine Microsecond Molecular Dynamics Simulations with AMBER on GPUs. 1. Generalized Born. J. Chem. Theory Comput. 8: 1542–1555.

Hansen E. E., Lozupone C. A., Rey F. E., Wu M., Guruge J. L., Narra A., Goodfellow J., Zaneveld J. R., McDonald D. T., Goodrich J. A., Heath A. C., Knight R., Gordon J. I., 2011 Pan-genome of the dominant human gut-associated archaeon, Methanobrevibacter smithii, studied in twins. Proceedings of the National Academy of Sciences of the United States of America 108: 4599–4606.

Heinicke I., Mller J., Pittelkow M., Klein A., 2004 Mutational analysis of genes encoding chromatin proteins in the archaeon Methanococcus voltae indicates their involvement in the regulation of gene expression. Mol Genet Genomics 272.

Henikoff S., Smith M. M., 2015 Histone variants and epigenetics. Cold Spring Harbor Perspectives in Biology 7: a019364.

Henneman B., van Emmerik C., van Ingen H., Dame R. T., 2018 Structure and function of archaeal histones. PLoS Genet 14: e1007582.

Higashibata H., Siddiqui M. A., Takagi M., Imanaka T., Fujiwara S., 2003 Surface histidine residue of archaeal histone affects DNA compaction and thermostability. FEMS Microbiology Letters 224: 17–22.

Jäger D., Förstner K. U., Sharma C. M., Santangelo T. J., Reeve J. N., 2014 Primary transcriptome map of the hyperthermophilic archaeon Thermococcus kodakarensis. BMC Genomics 15: 684.

Kazutaka Katoh D. M. S., 2013 MAFFT Multiple Sequence Alignment Software Version 7: Improvements in Performance and Usability. Mol Biol Evol 30: 772–780.

Kozlov A. M., Darriba D., Flouri T., Morel B., Stamatakis A., 2019 RAxML-NG: a fast, scalable and user-friendly tool for maximum likelihood phylogenetic inference. Bioinformatics 35: 4453–4455.

Le Grand S., Götz A. W., Walker R. C., 2013 SPFP: Speed without compromise—A mixed precision model for GPU accelerated molecular dynamics simulations. Computer Physics Communications 184: 374–380.

Letunic I., Bork P., 2019 Interactive Tree Of Life (iTOL) v4: recent updates and new developments. Nucleic Acids Research 47: W256–W259.

Luger K., Mäder A. W., Richmond R. K., Sargent D. F., Richmond T. J., 1997 Crystal structure of the nucleosome core particle at 2.8 Å resolution. Nature 389: 251–260.

Maier J. A., Martinez C., Kasavajhala K., Wickstrom L., Hauser K. E., Simmerling C., 2015 ff14SB: Improving the Accuracy of Protein Side Chain and Backbone Parameters from ff99SB. J. Chem. Theory Comput. 11: 3696–3713.

Malik H. S., Henikoff S., 2003 Phylogenomics of the nucleosome. Nat Struct Biol 10: 882–891.

Marc F., Sandman K., Lurz R., Reeve J. N., 2002 Archaeal histone tetramerization determines DNA affinity and the direction of DNA supercoiling. The Journal of Biological Chemistry 277: 30879–30886.

Mattiroli F., Bhattacharyya S., Dyer P. N., White A. E., Sandman K., Burkhart B. W., Byrne K. R., Lee T., Ahn N. G., Santangelo T. J., Reeve J. N., Luger K., 2017 Structure of histone-based chromatin in Archaea. Science 357: 609–612.

Maze I., Noh K.-M., Soshnev A. A., Allis C. D., 2014 Every amino acid matters: essential contributions of histone variants to mammalian development and disease. Nat. Rev. Genet. 15: 259–271.

Miller B. R. III, McGee T. D. Jr., Swails J. M., Homeyer N., Gohlke H., Roitberg A. E., 2012 MMPBSA.py: An Efficient Program for End-State Free Energy Calculations. J. Chem. Theory Comput. 8: 3314–3321.

Minh B. Q., Schmidt H. A., Chernomor O., Schrempf D., Woodhams M. D., Haeseler von A., Lanfear R., 2020 IQ-TREE 2: New models and efficient methods for phylogenetic inference in the genomic era. Mol Biol Evol 46: W537.

Mitchell A. L., Attwood T. K., Babbitt P. C., Blum M., Bork P., Bridge A., Brown S. D., Chang H.-Y., El-Gebali S., Fraser M. I., Gough J., Haft D. R., Huang H., Letunic I., Lopez R., Luciani A., Madeira F., Marchler-Bauer A., Mi H., Natale D. A., Necci M., Nuka G., Orengo C., Pandurangan A. P., Paysan-Lafosse T., Pesseat S., Potter S. C., Qureshi M. A., Rawlings N. D., Redaschi N., Richardson L. J., Rivoire C., Salazar G. A., Sangrador-Vegas A., Sigrist C. J. A., Sillitoe I., Sutton G. G., Thanki N., Thomas P. D., Tosatto S. C. E., Yong S.-Y., Finn R. D., 2019 InterPro in 2019: improving coverage, classification and access to protein sequence annotations. Nucleic Acids Research 47: D351–D360.

Nacev B. A., Feng L., Bagert J. D., Lemiesz A. E., Gao J., Soshnev A. A., Kundra R., Schultz N., Muir T. W., Allis C. D., 2019 The expanding landscape of “oncohistone” mutations in human cancers. Nature 567: 473–478.

Piquet S., Le Parc F., Bai S.-K., Chevallier O., Adam S., Polo S. E., 2018 The Histone Chaperone FACT Coordinates H2A.X-Dependent Signaling and Repair of DNA Damage. Molecular Cell 72: 888–901.e7.

Postberg J., Forcob S., Chang W.-J., Lipps H. J., 2010 The evolutionary history of histone H3 suggests a deep eukaryotic root of chromatin modifying mechanisms. BMC Evol. Biol. 10: 259.

Reeve J. N., Bailey K. A., Li W. T., Marc F., Sandman K., Soares D. J., 2004 Archaeal histones: structures, stability and DNA binding. Biochem. Soc. Trans. 32: 227–230.

Rojec M., Hocher A., Stevens K. M., Merkenschlager M., Warnecke T., 2019 Chromatinization of Escherichia coli with archaeal histones. eLife 8: 2407.

Salomon-Ferrer R., Götz A. W., Poole D., Le Grand S., Walker R. C., 2013 Routine Microsecond Molecular Dynamics Simulations with AMBER on GPUs. 2. Explicit Solvent Particle Mesh Ewald. . Chem. Theory Comput. 9: 3878–3888.

Sandman K., Grayling R. A., Dobrinski B., Lurz R., Reeve J. N., 1994 Growth-phase-dependent synthesis of histones in the archaeon Methanothermus fervidus. Proceedings of the National Academy of Sciences of the United States of America 91: 12624–12628.

Schymkowitz J., Borg J., Stricher F., Nys R., Rousseau F., Serrano L., 2005 The FoldX web server: an online force field. Nucleic Acids Research 33: W382–W388.

Shevchenko A., Tomas H., Havlis J., Olsen J. V., Mann M., 2006 In-gel digestion for mass spectrometric characterization of proteins and proteomes. Nat Protoc 1: 2856–2860.

Soares D. J., Marc F., Reeve J. N., 2003 Conserved Eukaryotic Histone-Fold Residues Substituted into an Archaeal Histone Increase DNA Affinity but Reduce Complex Flexibility. Journal of Bacteriology 185: 3453–3457.

Soares D. J., Sandman K., Reeve J. N., 2000 Mutational analysis of archaeal histone-DNA interactions. Journal of Molecular Biology 297: 39–47.

Talbert P. B., Henikoff S., 2010 Histone variants--ancient wrap artists of the epigenome. Nat Rev Mol Cell Biol 11: 264–275.

Tokura M., Ohkuma M., Kudo T., 2000 Molecular phylogeny of methanogens associated with flagellated protists in the gut and with the gut epithelium of termites. FEMS Microbiol. Ecol. 33: 233–240.

Williams T. A., Cox C. J., Foster P. G., Szöllősi G. J., Embley T. M., 2020 Phylogenomics provides robust support for a two-domains tree of life. Nat Ecol Evol 4: 138–147.

Wolfe J. M., Fournier G. P., 2018 Horizontal gene transfer constrains the timing of methanogen evolution. Nat Ecol Evol 2: 897–903.

Wollmann H., Stroud H., Yelagandula R., Tarutani Y., Jiang D., Jing L., Jamge B., Takeuchi H., Holec S., Nie X., Kakutani T., Jacobsen S. E., Berger F., 2017 The histone H3 variant H3.3 regulates gene body DNA methylation in Arabidopsis thaliana. Genome Biol. 18: 1–10.

